# Directed evolution of the Fe-nitrogenase for CO_2_ reduction to hydrocarbons

**DOI:** 10.64898/2026.07.08.737278

**Authors:** Niels N. Oehlmann, Frederik V. Schmidt, Jing Chen, Simone Prinz, Jan Zarzycki, Peter Claus, Jörg Kahnt, Tobias J. Erb, Johannes G. Rebelein

## Abstract

The iron (Fe) nitrogenase drives bacterial methane (CH_4_) formation by converting carbon dioxide (CO_2_) to CH_4_ in a single enzymatic step. Enhancing the initial CH_4_ formation activity of Fe-nitrogenase and expanding the product spectrum to hydrocarbon chains could lead to a route for sustainable feedstock chemicals. Here, we performed the first directed evolution campaign on the Fe-nitrogenase aimed at optimizing the hydrocarbon production. We achieved an ∼8-fold increase in CH_4_ formation by Fe-nitrogenase expressing *Rhodobacter capsulatus* cultures in three rounds of site-saturation mutagenesis. The best performing mutant (F362M*^anfD^*, Y85F*^anfD^*, T360S*^anfD^*) extends the *in vivo* product spectrum of the nitrogenase to ethane (C_2_H_6_) and exhibits 6-fold higher rates for CO production *in vitro*, whereas the formation of the undesirable byproduct formate was abolished. Electron microscopy–based structural analysis identified a methionine and water potentially stabilizing the transition state and fine-tuning the CO_2_ reduction mechanism and activity.

---

The sun is the primary energy source of planet earth.^1^ However, solar energy availability and energy demand fluctuate requiring energy storage solutions. Light energy can be converted to chemical fuels which are easily stored and distributed.^2,3^ The reduction of carbon dioxide (CO_2_) to hydrocarbons is attractive, because their distribution and storage infrastructure is available on large scale. Moreover, hydrocarbons are industrial feedstock chemicals, which replace fossil fuels and can contribute to carbon management and climate change mitigation strategies.^4–6^ The reduction of CO_2_ to CH_4_ is thermodynamically accessible in water.^7^ However, overpotentials and competing reaction pathways continue to limit efficiency and product selectivity. Therefore, improved catalysts for CH_4_ formation are desirable. Gas-processing metalloenzymes combine electron and proton delivery with defined active sites to achieve high selectivity and turnover.^8,9^ Moreover, these enzymes are easily produced in microorganism and can be further optimized by enzyme engineering.

The metalloenzyme nitrogenase is produced by prokaryotes, so called diazotrophs, and allows them to use molecular nitrogen (N_2_) as their nitrogen source. Three *bona fide* nitrogenase isozymes exist in nature (Fig. 1A). The common molybdenum (Mo)-nitrogenase is found in all diazotrophs and has the highest activity for nitrogen fixation.^10^ The alternative iron (Fe) and vanadium (V) nitrogenases function as fail-safe enzymes under Mo starvation.^11^ The Fe-nitrogenase is the only known natural enzyme that catalyses the direct reduction of CO_2_ to CH_4_ making it an interesting target for enzyme engineering.^12^ It consists of a reductase component (AnfH_2_, encoded by the *anfH* gene) and a catalytic component [Anf(DGK)_2_ encoded by the *anfDGK* gene, Fig. 1B].^13^ The reductase component AnfH_2_ is a homodimer with a subunit bridging [Fe_4_S_4_]-cluster and two ATP biding sites, allowing for ATP-dependent electron transfer to the catalytic component.^14,15^ Anf(DGK)_2_ is a heterohexamer and hosts the an electron relay, the [Fe_8_S_7_] P-cluster, and the active site cofactor, an [Fe_8_S_9_C-(*R*)-homocitrate] cluster, also termed iron-iron cofactor (FeFeco). During turnover, the S2B sulphide is mobilized, which exposes the Fe(2) and Fe(6) interface for substrate binding.^16^ This binding site is confined by the aliphatic residues V57*^anfD^* and F362*^anfD^.* In addition, the hydrogen-bond donors H180*^anfD^* and Q176*^anfD^*face the binding site and can interact with reduction intermediates.^15^ Turnover of nitrogenases is driven by the reductase component cycle (Fig. 1C). In this cycle, the reductase component is reduced by ferredoxins, transiently associates with the catalytic component, and donates a single electron to the P-cluster which shuttles the electrons to FeFeco to drive substrate reduction.^10,17,18^

**Fig. 1.**
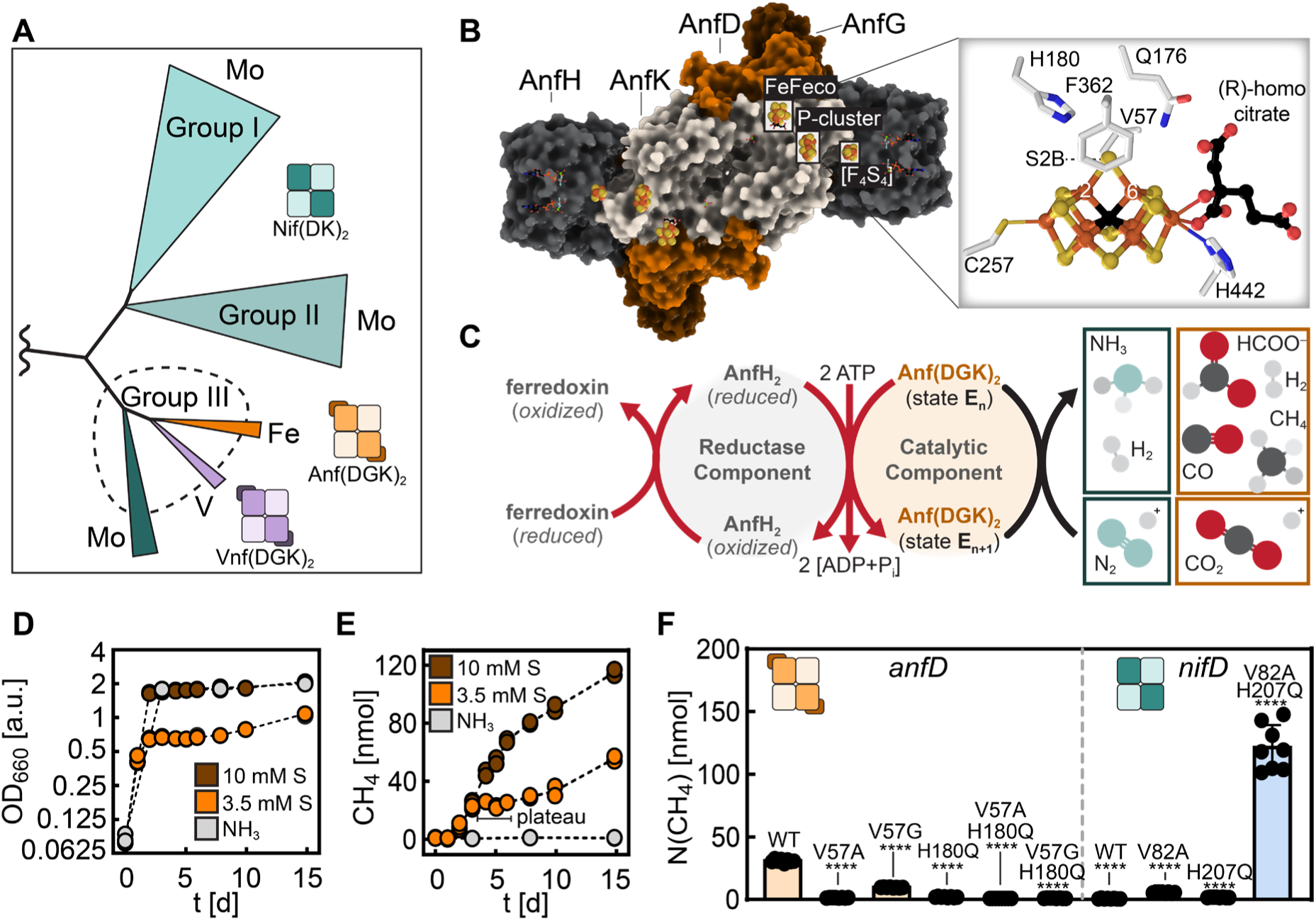
Methane formation by the Fe-nitrogenase. (A) Simplified scheme of the *bona fide* nitrogenase phylogenetic tree. The tree is based on the maximum-likelihood phylogenetic tree by Garcia *et al*.^35^ using the nomenclature by Raymond *et al*.^36^ Group I includes canonical Mo-nitrogenases from cyanobacteria and proteobacteria; Group II comprises Mo-nitrogenase homologs found in obligate anaerobes; and Group III encompasses the alternative V-and Fe-dependent nitrogenases, along with closely related Mo-nitrogenase homologs. (B) Structure of the Fe-nitrogenase (left, PDB: 8OIE) and close-up of the active site cofactor (right, FeFeco) highlighting the active site. (C) Reductase component cycle of the Fe-nitrogenase. Ferredoxins deliver electrons to the reductase component AnfH_2_. Subsequently, two ATP molecules bind to AnfH_2_, which interacts with the catalytic component Anf(HDGK)_2_ for electron transfer from the [Fe_4_S_4_]-cluster via the [Fe_8_S_7_] P-cluster to the FeFeco. The FeFeco is the site of small molecule reduction. Besides the reduction of N_2_ to NH_3_, the Fe-nitrogenase can convert CO_2_ to HCOO^-^, CO and CH_4_. Growth (D) or formation of CH_4_ (E) by Fe-nitrogenase expressing *R. capsulatus* cultures over time using NH_3_ or serine (S) as nitrogen source. Each dot represents an individual culture. n = 2 independent experiments per time point. (F) Amount of CH_4_ formed by Fe-nitrogenase and Mo-nitrogenase active site mutants after 4 d of cultivation. Each dot represents an individual culture. n = 8 independent experiments. Statistical analysis was performed using unpaired two-tailed Welch’s t-tests against the WT: ****P ≤ 0.0001.

Nitrogenases are promiscuous enzymes.^10,18,19^ The V nitrogenases reduces carbon monoxide (CO) to CH_4_ and short-chain hydrocarbons, mainly forming ethylene (C_2_H_4_) a reaction that mimics the Fischer-Tropsch process under biological conditions.^11,20–23^ The Fe-nitrogenase is best in reducing CO_2_ mainly forming formate (HCOO^-^) and CH_4_.^12,24–27^ Both, HCOO^-^ and CH_4_ may serve as nutrients for other microbes shaping microbial communities and are important industrial feedstock chemicals.^28,29^ Nitrogenase-mediated CO_2_ reduction has therefore attracted interest for its relevance and potential in a sustainable bioeconomy.^30^ Here, we sought to enhance the CO_2_-reduction activity of the Fe-nitrogenase toward hydrocarbon formation.

Previously, site-directed mutagenesis on the *Azotobacter vinelandii* nitrogenase was used to investigate its mechanism and to alter the CO_2_ activity.^9^ An active-site double mutant (V70A*^nifD^*,H195Q*^nifD^*, *nifD*: Mo- nitrogenase structural gene) of the Mo-nitrogenase described by Seefeldt and co-workers could reduce CO_2_ to CH_4_.^12,31,32^ Moreover, the V70A*^nifD^* mutant showed increased electron flux towards HCOO^-^formation.^27^ However, the difficult genetic manipulation of *A. vinelandii* and the lack of expression plasmids aggravate the work with this model organism. Thus, only a limited number of nitrogenase mutants were screened.^33,34^ Using our recently established plasmid-based nitrogenase expression system for *Rhodobacter capsulatus*,^14^ we performed the first directed evolution campaign on the nitrogenase catalytic component to increase the CH_4_ formation of the Fe-nitrogenase. First, we developed a screening method for the quantification of nitrogenase-derived CH_4_ and developed a workflow to screen site-saturation mutagenesis libraries in *R. capsulatus*. By performing three rounds of directed evolution, we obtained the Fe-nitrogenase variant V3 (F362M*^anfD^*, Y85F*^anfD^*, T360S*^anfD^*) with an 8-fold increase *in vivo* CH_4_ formation. Moreover, *in vitro* analysis revealed that V3 forms CH_4_ and CO at 2.6-fold and 6-fold higher rates, while abolishing the detrimental HCOO^-^ formation activity. Structural characterization of V3 by electron microscopy showed a methionine and water molecules in proximity to the FeFeco, which we propose to fine-tune the mechanism to favor formation of CH_4_ over HCOO^-^. These experimental results establish directed evolution for nitrogenase and underline the potential of the Fe-nitrogenase as a CO_2_ reduction catalyst for a circular bioeconomy.

## Directed evolution of nitrogenase

First, we established an *in vivo* readout for CH_4_ formed by the Fe-nitrogenase in *R. capsulatus* strain B10S.^37^ The chromosomal Mo- and Fe-nitrogenase were removed. In addition, the Mo transporter ModABC was deleted to promote the Fe-nitrogenase expression and the *draTG* gene (reductase component ribosyltransferase) was removed to prevent deactivation of AnfH_2_. The Fe-nitrogenase was reconstituted on a plasmid using the native promotor to obtain the expression strain MM0167 (B10S *ΔnifD::SpR ΔmodABC ΔanfHDGK::GmR ΔdraTG* carrying the Fe-nitrogenase expression plasmid pMM0128).

Nitrogenase *in vivo* CH_4_ activity is dependent on several factors. It can vary with nitrogenase expression levels, growth phase, metabolic activity, light intensity and CO₂ concentration, among other factors.^12^ Therefore, we decided to assess the *in vivo* nitrogenase activity with CH_4_ quantification after cultures have reached a plateau in CH_4_ formation to average out kinetic differences. For the CH_4_ assay, we conducted growth of the *R. capsulatus* strains in gas chromatography (GC) vials illuminated by LED panels at 30 °C compatible with CH_4_ quantification via GC-FID (Fig. 1D,E). Serine is used as an alternative nitrogen source, as NH_3_ represses the Fe-nitrogenase. Following Schneider *et al.*,^38^ we compared limiting serine concentrations (3.5 mM) to standard RCV media (10 mM serine). The CH_4_ production was monitored over time to determine a suitable timepoint for the CH_4_ quantification. As expected, the NH_4_ ^+^ supplemented control showed no CH_4_ formation. Cultures supplemented with 10 mM serine reached the same optical density at 660 nm (*OD*_660_) as the NH_4_ ^+^ supplemented control (Fig. 1D). CH_4_ formation was first detected after 2 days, when the cultures reached stationary phase. Afterwards, the formation of CH_4_ proceeded until the end of the experiment (15 d). In contrast, reducing the serine concentration to 3.5 mM limits culture growth and changes the CH_4_ time profile. CH_4_ formation reaches a plateau between days 4 and 6, providing a suitable time window for the CH_4_ quantification. Taken together, we established cultivation conditions for *in vivo* Fe-nitrogenase activity assays and CH_4_ quantification.

To test our set-up, we determined the CH_4_ formation of Fe and Mo-nitrogenase wild type (WT) as well as mutants constructed based on the best known CH_4_ forming Mo-nitrogenase double mutant V82A*^nifD^*, H207Q*^nifD^*(corresponding to V70A*^nifD^* and H195Q*^nifD^* in the *A. vinelandii nifD* numbering; Fig. 1E).^31,32^ The Fe-nitrogenase WT produced CH_4_ (∼30 nmol). Mutating V57*^anfD^* to glycine or alanine either reduced or prevented CH_4_ formation for the Fe-nitrogenase. Similarly, the H180Q*^anfD^* mutant alone or in combination with the V57G*^anfD^* or V57A*^anfD^* showed no CH_4_ formation confirming the surprising results of the *Rhodopseudomonas palustris* Fe-nitrogenase mutants constructed by Zheng *et al.*^12^ In contrast, the Mo-nitrogenase double mutant (V82*^nifD^*, H207Q*^nifD^*) showed a 4-fold higher final CH_4_ amount (∼120 nmol) compared to the Fe-nitrogenase WT. In summary, we established an *in vivo* screen for CH_4_ formation and demonstrated its effectiveness for variant identification using known mutants. Moreover, the results suggest that Mo- and Fe-nitrogenase differ significantly in their CO_2_ reduction, as the active site mutations show distinct effects on CH_4_ formation.

Next, we set up a directed evolution workflow for the Fe-nitrogenase based on the *in vivo* CH_4_ assay (Extended data Fig. 1 for detailed description). Initial attempts to introduce site-saturation mutagenesis libraries in the broad host range plasmid pMM0128 (pBBR1 replicon) suffered from low cloning efficiencies. Therefore, we developed a two-step strategy relying on a high copy number plasmid (ColE1 replicon) to generate mutant libraries of *anfD* by using degenerate NNK codons.^39^ Afterwards, the *anfD* libraries were transferred via golden gate cloning into pMM0128. The library was then conjugated into *R. capsulatus* and cultivated in 96 well microtiter plates to generate precultures for the CH_4_ assays. The workflow requires 18 days from the primer design to the final CH_4_ quantification (library cloning: 4 d, conjugation into *R. capsulatus*: 4 d, preculture cultivation: 6 d, CH_4_ assay cultivation: 4 d) allowing to screen thousands of mutants per month.

Three cycles of directed evolution were conducted (Fig. 2A,B). Initially, we focused on residues close to the FeFeco known for their involvement in nitrogenase catalysis (*i.e.* V57*^anfD^*, H180*^anfD^*, K83*^anfD^*, Q176*^anfD^*).^9^ After these libraries did not provide improved variants, we screened a combinatorial library (based on the “smart” library design) of the two glycine residues G336*^anfD^* and G337*^anfD^* that were shown to bind N_2_ reduction intermediates.^40^ Again, no improved CH_4_ variant was found. Following this setback, we screened 21 site-saturation mutagenesis libraries and another combinatorial library (V57*^anfD^* and H180*^anfD^*) before we identified a first improved variant for CH_4_ formation targeting the F362*^anfD^* residue. The F362M*^anfD^* mutation resulted in ∼4-fold more CH_4_ formation compared to the WT. After the improved activity of the F362M*^anfD^* mutant (V1) was confirmed, by retransforming the plasmid into the base strain to exclude a genomic mutation causing this improvement, we locked the mutation. In cycle 2, we screened eleven site-saturation mutagenesis libraries and locked the Y85F*^anfD^* mutation (V2) with slightly increased *in vivo* CH_4_ yields.

**Fig. 2.**
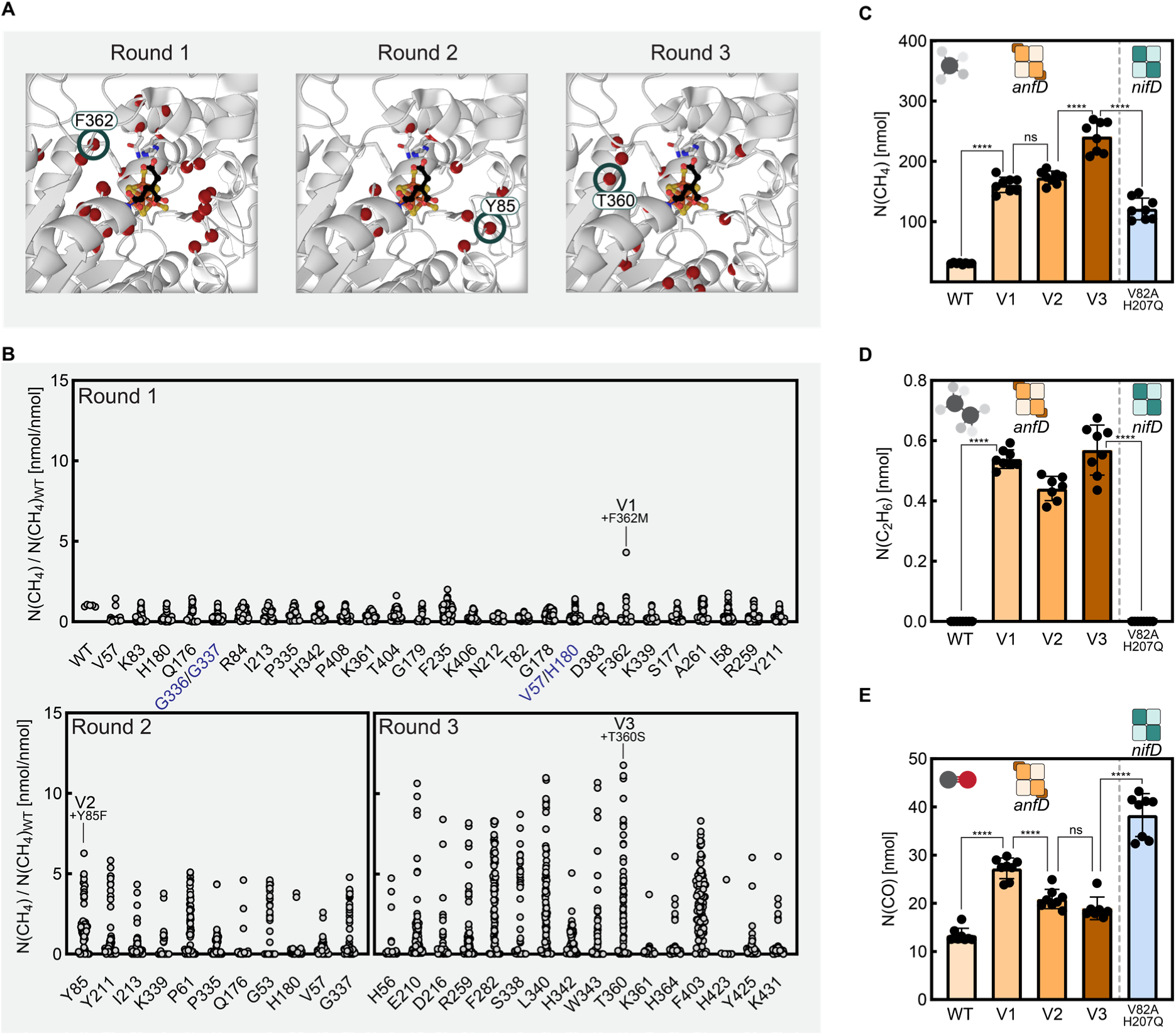
Directed evolution of the Fe-nitrogenase for CH_4_ formation. A plasmid-based site-saturation mutagenesis (SSM) library of the nitrogenase *anfD* gene is generated by amplification with degenerate oligonucleotides and subcloned into a broad host range plasmid by Golden Gate cloning (Extended Data Figure 1). The library is conjugated into the *R. capsulatus* Fe-nitrogenase expression strain MM167 and single colonies are cultivated in sealed headspace sampling gas chromatography vials (4 mL RCV media in 20 mL vials) under anaerobic and phototrophic conditions (30 °C, 8% CO_2_ in argon atmosphere, 4 d). The final CH_4_ and C_2_H_6_ amount is quantified by headspace sampling GC. The best performing variants were sequenced and a mutation was locked for the next round of evolution after a verification *in vivo* assay using more replicates. (A) Structure of the FeFeco environment in *anfD* (PDB: 8OIE). Targeted residues of the directed evolution rounds are highlighted in red. Green circles indicate the site of the locked mutation. (B) Final amount of CH_4_ formed in the *in vivo* assays during the directed evolution normalized to the WT Fe-nitrogenase CH_4_ formation. Dots represent individual CH_4_ amounts of one *R. capsulatus* culture. A single site SSM library based on NNK codon was evaluated with 96 cultures. The combinatorial SSM libraries (blue label) based on the “smart” codon design were evaluated with 1200 cultures. Final amount of CH_4_ (C), C_2_H_6_ (D) and CO (E) in *in vivo* CH_4_ assays of *R. capsulatus* cultures expressing nitrogenase mutants (30 °C, 8% CO_2_ in argon atmosphere, 4 d). Each dot represents an *in vivo* activity assay. n = 8 biological replicates. Statistical analysis was performed using unpaired two-tailed Welch’s t-tests: ****P ≤ 0.0001.

A third cycle identified mutation T360S*^anfD^* in proximity to the first F362M*^anfD^* mutation (V3), producing 2-fold more CH_4_ compared to the double mutant of the Mo-nitrogenase (V82A*^nifD^*, H2075Q*^nifD^*). It appears that mutant V2 (with F362M*^anfD^*Y85F*^anfD^*) overcame a local maximum and opened up the mutational landscape since several improved variants were identified. We confirmed the increased activity of V1-V3 by performing biological replicates in the original base strain, with V3 reaching a final amount of 241 nmol CH_4_ (Fig. 2C).

Excitingly, small amounts of C_2_H_6_ were produced by cultures expressing the variants (Fig. 2D). This product is not formed by cultures expressing the WT Fe-nitrogenase or Mo-nitrogenase mutants indicating that the Fe-nitrogenase variants have an expanded product profile. Furthermore, all variants show increased amounts of carbon monoxide (CO) in the culture headspace (Fig. 2E). CO is a known substrate of the Fe-nitrogenase for CH_4_ formation that could also be involved in the C-C coupling reaction.^22^

### *In vitro* activity analysis

To unveil the reasons for increased *in vivo* CH4 formation, we assessed the variants by *in vitro* activity assays. The Fe-nitrogenase variants were purified by affinity chromatography (Extended data Fig. 2). *In vitro* assays were conducted under 1.2 atm CO_2_ (Fig. 3A-B, E-F). The WT variant produces HCOO^−^ {49 ± 4 nmol × [nmol Anf(DGK)_2_ × min]^-1^} as the main product and lower amounts of H_2_ {32.8 ± 0.4 nmol × [nmol Anf(DGK)_2_ × min]^-1^} (Fig. 3A,B). In stark contrast, all variants show a ∼2-fold higher H_2_ formation rate {V3: 77 ± 3 nmol × [nmol Anf(DGK)_2_ × min]^-1^} as well as a as a 2.6-fold increased CH_4_ formation rate {V3: 0.081 ± 0.008 nmol × [nmol Anf(DGK)_2_ × min]^-1^ vs. WT: 0.0316 ± 0.0004 nmol × [nmol Anf(DGK)_2_ × min]^-1^ but a strongly reduced rate for HCOO^−^ formation ({V3: 1.0 ± 0.4 nmol × [nmol Anf(DGK)_2_ × min]^-1^}; Fig. 3A,B,F). Thus, our directed evolution likely selected for the formation of CH_4_ and H_2_ over HCOO^−^.

**Fig. 3.**
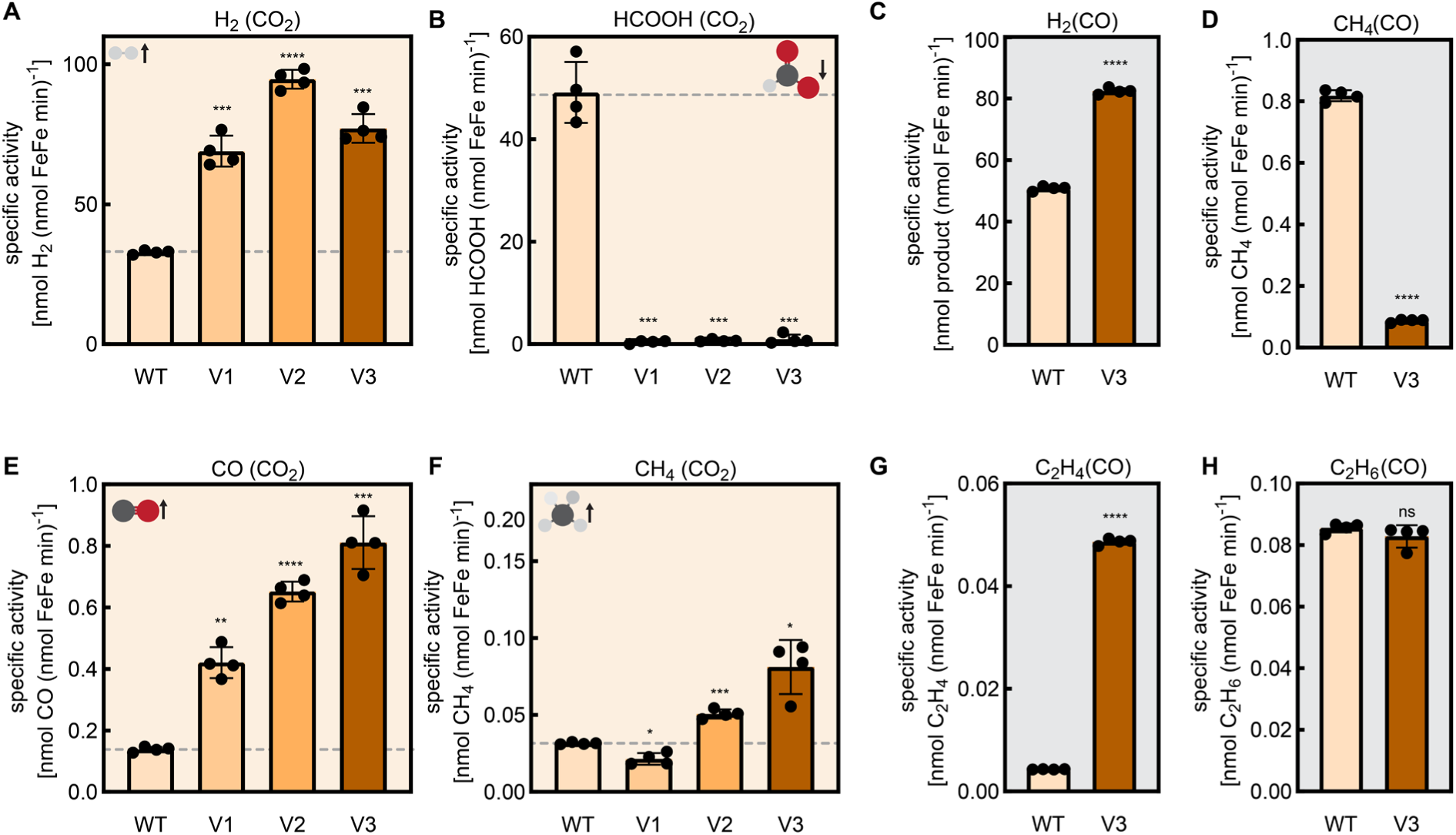
CO_2_ and CO reduction activity of the purified Fe-nitrogenase mutants. (A-B, E-F) Product formation rates for *in vitro* activity assays conducted under a 1.2-atm CO_2_ atmosphere using 0.1 mg Anf(DGK)_2_ and 0.5 mg (20× molar excess) AnfH_2_. Plotted are specific activities for the formation of H_2_ (A), HCOO^−^ (B), CO (E) and CH_4_ (F). Each dot represents an activity assay. n = 4 independent experiments. (C-D, G-H) Product formation rates for *in vitro* activity assays conducted under 0.09 atm CO. Plotted are specific activities for the formation of H_2_ (C) and CH_4_ (D) and C_2_H_4_ (G) and C_2_H_6_ (H). Each dot represents an activity assay. n = 4 independent experiments. Statistical analysis was performed using unpaired two-tailed Welch’s t-tests against the WT: ns: not significant, *P ≤ 0.05, **P ≤ 0.01, ***P ≤ 0.0001, ****P ≤ 0.0001.

In addition, the directed evolution resulted in an ∼6-fold increase of the CO formation rate ({V3: 0.81 ± 0.04 nmol × [nmol Anf(DGK)_2_ × min]^-1^ vs. WT: 0.138 ± 0.004 nmol × [nmol Anf(DGK)_2_ × min]^-1^}, Fig. 3E). The formation of C–C-coupling products was however not observed in the CO_2_ reduction assays. Importantly, the total electron flux of the variants is similar to the WT (V3: 158 min^-1^ vs. WT: 165 min^-1^} demonstrating full activity of the reductase component cycle (Extended data Fig. 3). Overall, our results demonstrate that directed evolution enhanced *in vivo* CH_4_ formation by suppressing HCOO^−^ and increasing the rate for CO and CH_4_ formation. Moreover, the mutations did not interfere with the electron transport by the reductase component cycle.

To illuminate the formation of C_2_H_6_ by cultures expressing variant V3, we performed CO reduction assays. The assays were conducted at 0.09 atm CO in an argon atmosphere to maximize the CO reduction activity (Fig. 3C-D, G-H). While V3 has an increased rate for H_2_ formation under CO (Fig. 3C), the rate for CH_4_ formation under CO is 8-fold lower compared to the WT {V3: 0.087 ± 0.002 nmol × [nmol Anf(DGK)_2_ × min]^-1^ vs. WT: 0.819 ± 0.004 nmol × [nmol Anf(DGK)_2_ × min]^-1^}. However, V3 has an improved C–C-coupling activity. V3 produces ethylene (C_2_H_4_) at a rate of 0.0487 ± 0.0003 nmol × [nmol Anf(DGK)_2_ × min]^-1^, whereas C_2_H_4_ formation by the WT is negligible. Both V3 and the WT produce C_2_H_6_ at similar rates (0.0829 ± 0.0016 *vs.* 0.0856 ± 0.0006 nmol × [nmol Anf(DGK)_2_ × min]^-1^, respectively).

Besides CO_2_ and CO reduction activity, we investigated changes in the reduction of the natural substrate N_2_ (Extended Data Fig. 4) and acetylene (C_2_H_2_, Extended Data Fig. 5). To our surprise, the variants were still able to reduce N_2_ to NH_3_ indicating that the mutations did not interfere with the Lowe−Thorneley cycle including the formation of the E_4_ state (Extended Data Fig. 4A). This suggests that the capacity for N_2_ reduction and CH_4_ formation could be linked in Fe-nitrogenase. Nevertheless, the mutants were unable to support diazotrophic growth of *R. capsulatus* (Extended Data Fig. 4B), demonstrating that the WT nitrogenase is subjected to selective pressures beyond N_2_ reduction *in vivo*. The discrepancy between *in vitro* activity assays and *in vivo* activity is also illustrated by the C_2_H_2_ reduction assays. In *R. capsulatus* whole cell C_2_H_2_ reduction assays performed prior to nitrogenase purification, we observed increased C_2_H_6_ formation compared to the WT (Extended Data Fig. 5A-C). While the WT Fe-nitrogenase culture produced 2% C_2_H_6_ during the assay, variants V1, V2, and V3 produced 49%, 39%, and 30% C₂H₂, respectively. When we repeated the C_2_H_2_ reduction assay with purified nitrogenases (Extended Data Fig. 5D-F), V3 still exhibited higher activity for C_2_H_6_ formation compared to the WT (V3: 0.870 ± 0.012 nmol × [nmol Anf(DGK)_2_ × min]^-1^; WT: 0.453 ± 0.016 nmol × [nmol Anf(DGK)_2_ × min]^-1^) but the percentage of C_2_H_6_ was lowered to 15% showcasing the differences of *in vivo* and *in vitro* conditions. Nevertheless, the C_2_H_2_ reduction assays demonstrate that the mutations have a strong influence on the product profile but maintain high overall activity. In conclusion, directed evolution preserved N_2_ fixation activity, indicating a coupling to the CH_4_ formation activity. In addition, we observed a shift in C_2_H_2_ reduction products from C_2_H_4_ to C_2_H_6_. Both activities, C_2_H_2_ and N_2_ reduction were deviating *in vivo* from the *in vitro* observations underscoring that *in vitro* assays may not fully reflect cellular conditions, particularly regarding electron availability and transport. The *in vitro* assays were performed at high electron flux conditions with a 20-fold excess of reductase to catalytic component, while in *in vivo* only a 5-fold excess is expected.^41^

To conclude, we conducted a comprehensive *in vitro* activity analysis for the Fe-nitrogenase variants and find the CH_4_ formation by *R. capsulatus* cultures in agreement with changes in product profile and activity of the variants.

### Structural analysis of V3

We prepared the Anf(DGK)_2_(H_2_)_2_ complex of V3 by trapping the ADP-bound reductase component on the Anf(DGK)_2_ core with AlF_3_ and performed cryo-EM structural analysis (Table 1). Using the cryo-EM map (global resolution 2.04 Å), we built a model of V3 (Fig. 4A). All nitrogenase cofactors were intact and well resolved in accordance with the high electron flux of V3 (Fig. 4B-D). The structural features of the [Fe_4_S_4_] cluster and the P-cluster resemble the WT and are not affected by the V3 mutations.

**Fig. 4.**
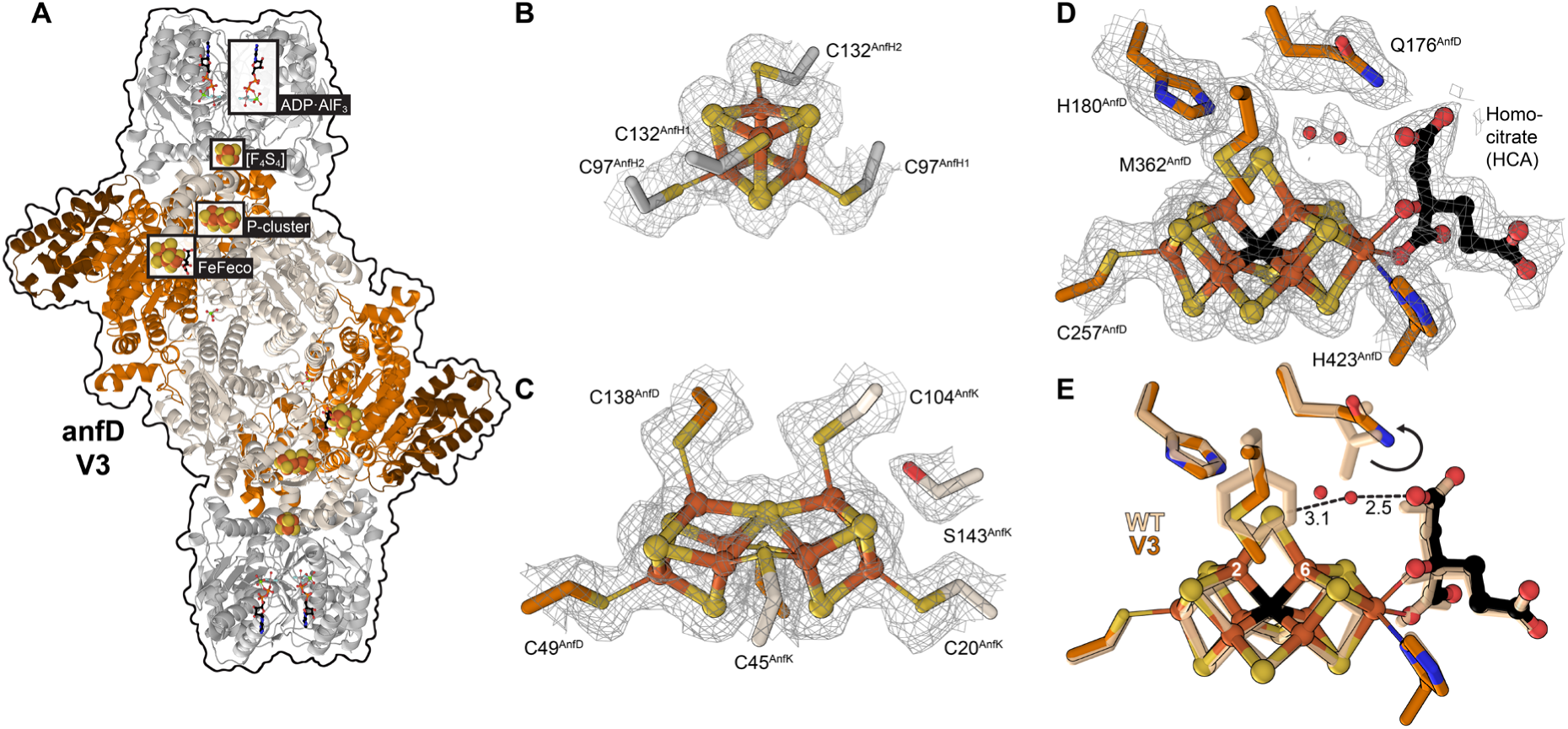
Structure of the Fe-nitrogenase mutant V3. (A) Model of the Fe-nitrogenase mutant V3. Cofactors are highlighted by grey boxes. Close-up views of the [Fe_4_S_4_] cluster (B), the P-cluster (C) and the FeFeco (D) overlaid with a mash representation of the cryo-EM density map. (E) Overlay of the active site (FeFeco, Q176^anfD^, H180^anfD^, M/F362^anfD^) of V3 (orange) and the WT (wheat) Fe-nitrogenase.

**Table 1.**
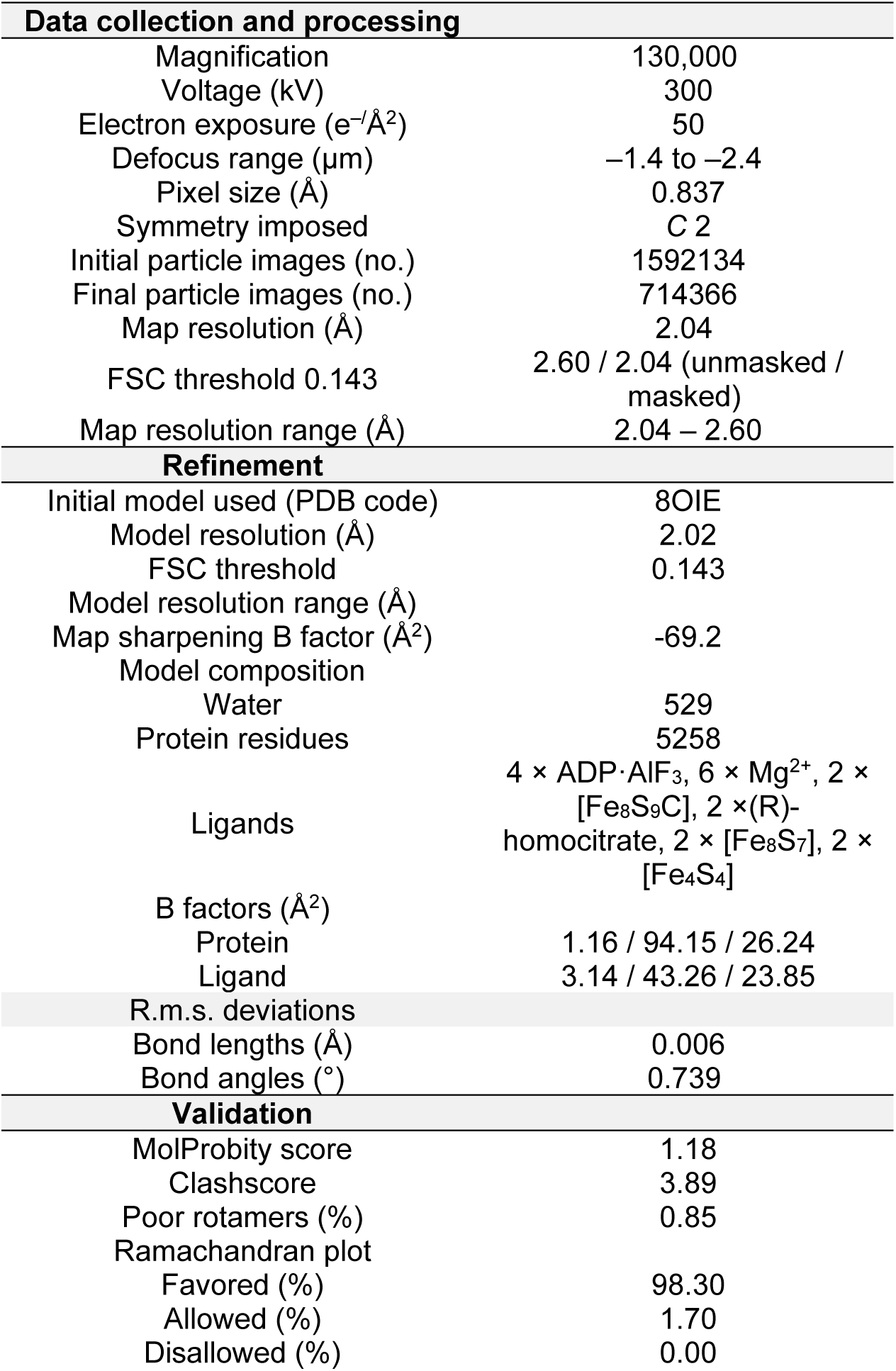
Cryo-EM data collection, refinement, and validation statistics.

Next, we analysed the structure for features that could explain the activity changes. The largest CH_4_ activity increase of V3 was caused by the F362M*^anfD^* and T360S*^anfD^*mutations. They are conservative mutations close to the FeFeco. Interestingly, both mutations reduce the sidechain volume [Mean volumes (*V*) of residues buried in protein interiors: *V*(F): 193.5 Å^3^; *V(*M): 167.7 Å^3^; *ΔV*(F→M): −25.8 Å^3^; *V*(T): 120.0 Å^3^; *V*(S): 94.2 Å^3^; *ΔV*(T→S): −25.8 Å^3^].^42^ This opens up enough volume inside the protein for two water (H_2_O) molecules (Mean *V* of H_2_O buried in protein interiors: 24.5 Å^3^).^43^ Indeed, our cryo-EM map had two unassigned globular densities near the FeFeco that we modelled as H_2_O (Fig. 4D). The H_2_O molecules form hydrogen bonds with the S2B sulphide and the (*R*)-homocitrate ligand of the FeFeco. By comparing the active site structure of V3 to the WT, we observed that the H_2_O molecules displaced residue Q176*^anfD^* into the resting state position described by Einsle and co-workers (Fig. 4E). Q176*^anfD^* has been proposed to interact with the S2B sulphide and reaction intermediates, while (*R*)-homocitrate is involved in the proton delivery. Thus, the H_2_O molecules interact with key features of the nitrogenase mechanism.

Mutation F362M*^anfD^* also introduces a Lewis basic sulphur atom into the second coordination sphere of the FeFeco. C-S-bonds can be substrates for nitrogenase-like enzymes. For example, the nitrogenase-like enzyme methylthio-alkane reductase cleaves the C-S-bond of methylthioethanol. To exclude C-S-bond cleavage of F362M*^anfD^*as a potential CH_4_ source, we performed mass spectrometry and peptide identification on V3 to identify modifications of F362M*^anfD^* (Extended Data Fig. 6). No modification was found. In summary, we obtained a 2.04 Å resolution map of the V3 Anf(DGK)_2_(H_2_)_2_ complex revealing F362M*^anfD^* and two H_2_O molecules, together they might alter the hydrogen bonding network at the FeFeco and influence the CO_2_ reduction mechanism.

## Discussion

Here, we established an *in vivo* CH_4_ activity assay for the Fe-nitrogenase, established a directed evolution workflow and performed three cycles of directed evolution. We screened 25 individual site-saturation mutagenesis libraries and two combinatorial libraries to identify V1 (Fig. 2C), indicating a low mutational flexibility of the Fe-nitrogenase. This can be explained by the intricate catalytic cycle involving the transfer of eight electron and protons for CH_4_ formation.^18,27^ Moreover, the electron transfer as well as the catalysis is determined by metalloclusters which cannot be mutated, thus limiting the mutational space. Taken together, we observe a high screening burden. Nevertheless, our method was successful in increasing CH_4_ formation, and we expect to find mutants with higher activity in the future.

The variant V3 exhibited remarkable activities *in vitro*. The formation of CO and CH_4_ was increased 6-fold and 2.6-fold, respectively. Simultaneously, the electron flux to HCOO^−^ has been shifted to the formation of H_2_ by V3. We previously showed that HCOO^−^ is secreted by Fe-nitrogenase expressing *R. capsulatus* cultures, rendering it a dead-end metabolite.^26^ In contrast, H_2_ can be recovered via uptake hydrogenases.^44^ Hence, supressing HCOO^−^ increases the available resources for CH_4_ formation. Furthermore, V3 shows an increased C-C-coupling activity, which combined with the increased formation of CO from CO_2_ is likely responsible for the formation of C_2_H_6_ *in vivo*, the first observation of C-C-coupling products from CO_2_ by a nitrogenase.

Finally, we performed cryo-EM structure analysis of V3 (Fig. 4). We observed a H_2_O molecule binding between the *R*-homocitrate and the S2B belt sulphide. In addition, F362M*^anfD^* introduces a Lewis base facing towards the FeFeco S2B position. Fig. 5A shows a mechanistic proposal for nitrogenase catalysed CO_2_ reduction based on the reaction pathways described by Khadka *et al.*^27^ and the mechanistic model proposed by Rohde *et al.*^45^ Nitrogenase catalysis proceeds via stepwise e^−^/H^+^ transfer to the FeFeco, which increases the E_n_ state of the cofactor (Lowe–Thorneley scheme).^17^ After the FeFeco reaches the E_2_ state, the S2B belt sulphide is replaced by an Fe-bound hydride. CO_2_ can react with the E_2_ state following two pathways. In the direct hydride transfer (DHT), the hydride undergoes bond formation with the carbon atom of CO_2_ forming HCOO^−^. In the associative pathway (AP), the CO_2_ molecule is first activated by the Fe atom. Afterwards, a proton transfer to the CO_2_ O-atom results in the cleavage of H_2_O and the formation of CO. The metal bound CO can be further reduced to CH_4_ or hydrocarbons via CO insertion following the mechanism described by Rohde *et al.*^45^

**Fig. 5.**
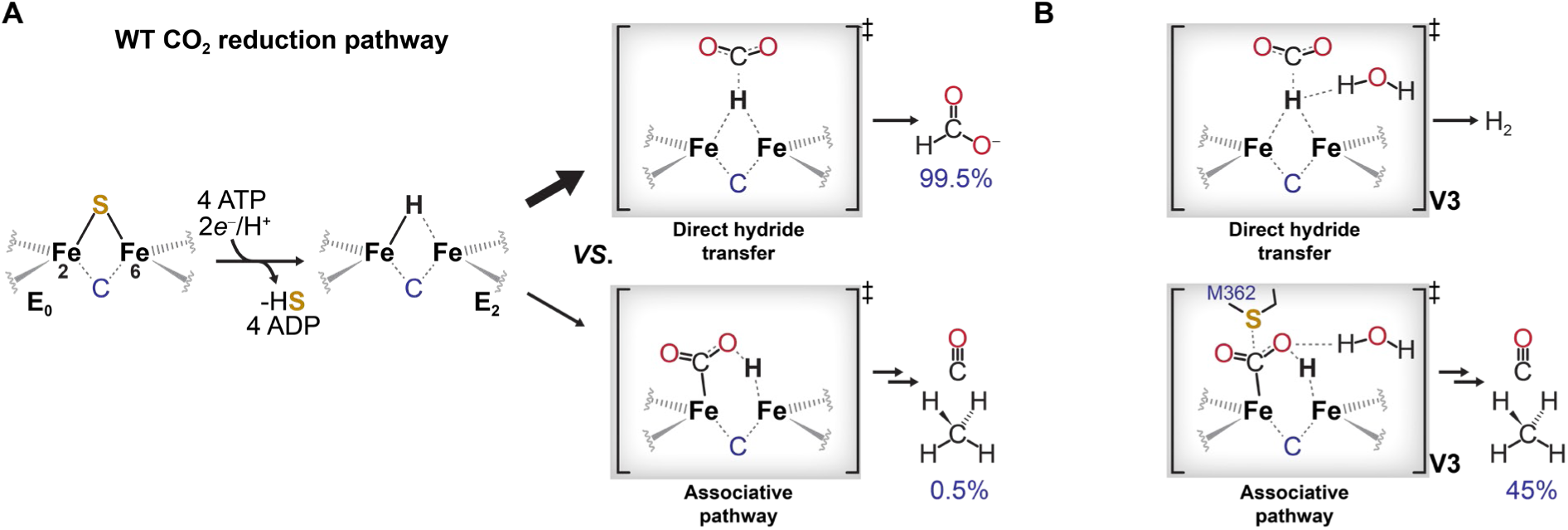
Mechanistic proposal of CO_2_ reduction by nitrogenases. (A) Mechanistic model based on the model described by Khadka *et al.* and Rhode *et al.* The ground state E_0_ of the FeFeco is converted to the E_2_ state by the transfer of two electrons to the FeFeco resulting in a loss of the S2B belt sulfide. The E_2_ state reacts with CO_2_ following two pathways. CO_2_ is converted to HCOO^−^ by a direct hydride transfer (DHT) from the FeFeco to the CO_2_ carbon atom. Alternatively, the CO_2_ molecule can be activated by the Fe atom followed by the protonation of one CO_2_ oxygen atom in the associative pathway (AP). The associative pathway results in the cleavage of water and the formation of CO and CH_4_. The DHT is preferred by the WT Fe-nitrogenase resulting in 99.5% HCOO^−^ yield (B) Proposed influence of the point mutations in V3 on the transition states of the DHT and AP transition states. The FeFeco bound proton reacts as a hydride in the DHT transition state. The water molecule in V3 found in close proximity to the FeFeco could function as an alternative acceptor for the hydride (instead of CO_2_) resulting in the formation of H_2_ (instead of HCOO^−^). The FeFeco bound proton has protic character in the transition state of the AP, preventing the formation H_2_ by the water molecule. Instead, the water molecule could facilitate the O-protonation. Furthermore, the Lewis basic methionine M362 could support the CO_2_ activation to facilitate the C−Fe bond formation.

The Fe-nitrogenase WT favours DHT over AP, with 99.5% of the CO_2_ directed electron flux ending in HCOO⁻ formation. Fig. 5B shows the potential impact of the H_2_O molecules in V3 on the transition state of the DHT and AP. The Fe bound hydrogen has hydridic character in the DHT transition state. The H_2_O can react with the hydride to liberate H_2_ preventing the hydride transfer to the CO_2_. This explains the redirection of electron flux from HCOO^−^ to H_2_ in V3.

In contrast, the metal bound hydrogen has protic character in AP transition state, which is unable to react with the H_2_O molecule. Moreover, H_2_O could facilitate the water cleavage for CO formation from the AP transition state by providing a second proton for O-protonation. The Lewis basic sulphur of F362M*^anfD^*could further help to activate the CO_2_ molecule analogues to the mechanism described by Ryde and co-workers for the formate dehydrogenase.^46^ In this mechanism, CO_2_ interacts with the cysteine ligand of the molybdenum cofactor, decreasing the O−C−O-bond angle and facilitating the nucleophilic attack on the carbon atom. This is in line with the increase of CO_2_ directed electron flux towards CO and CH_4_ formation in V3 (45 %). In conclusion, the structural characterization of V3 showed changes around the FeFeco. Based on these, we proposed a mechanistic model by which altered proton transfer at the FeFeco could favour H_2_, CO and CH_4_ formation and prevents the formation of HCOO^−^.

Overall, our work demonstrates that directed evolution is a powerful approach for uncovering nitrogenase functionalities, paving the way for mechanistic studies and future applications in nitrogenase-based CO₂ reduction.

### Materials and Methods

#### 1.1.1 Chemicals

Unless noted otherwise, all chemicals were purchased from Fisher Scientific Inc. (Waltham, USA), Carl Roth GmbH + Co. KG (Karlsruhe, Germany), Thermo Sigma-Aldrich (St. Louis, USA), or Tokyo Chemical Industry Deutschland GmbH (Eschborn, Germany) and were used directly without further purification. Gases were purchased from Air Liquide Deutschland GmbH (Düsseldorf, Germany)

#### 1.1.2 Strains, plasmids, and growth conditions

All strains used in this study are listed in Supplementary Table 1. *E. coli* is cultivated aerobically on Luria–Bertani (LB) media (10 g/L tryptone, 5 g/L yeast extract and 10 g/L NaCl dissolved in deionized water) agar plates at 37 °C. Liquid cultures are grown in 10 mL LB media in 100 mL Erlenmeyer flasks, shaking at 180 rpm at 37 °C*. R. capsulatus* is cultivated in *R. capsulatus* minimal medium RCV [containing 30 mM DL-malic acid, 0.8 mM MgSO_4_, 0.7 mM CaCl_2_, 0.05 mM sodium ethylenediamine-tetraacetic acid (Na_2_EDTA), 0.03 mM thiamine hydrochloric acid, 10 mM KH_2_PO_4_, 120 µM FeSO_4_, 45 µM B(OH)_3_, 9.5 µM MnSO_4_, 0.85 µM ZnSO_4_, 0.15 µM Cu(NO_3_)_2_ at a pH set to 6.8; supplemented with 10 mM or 3.5 mM serine as N-source] or on complex Peptone-Yeast medium (PY; containing 10 g/L peptone, 0.5 g/L yeast extract, after autoclaving 2 mM MgCl_2_, 2 mM CaCl_2_, 80 μM FeSO_4_ are added) agar plates. Cultivation is conducted under anaerobic and phototrophic growth conditions (illuminated by 6 x 60 W krypton light bulbs (Osram Licht AG, Munich, Germany) or custom-built LED panels (λ: 850 nm, 420 nm and 470 nm, ∼60 μmol photons m^-2^ s^-1^) at 30 °C adapted from Katzke *et al*.^47^ Agar plates of *R. capsulatus* are cultivated for 2 d in Oxoid™ 3.5 L anaerobic jars (Thermo Fisher Scientific Inc., Waltham, USA) anaerobised using Oxoid™ AnaeroGen™ 3.5L sachets (Thermo Fisher Scientific Inc., Waltham, USA). Liquid cultures of *R. capsulatus* strains are grown in 4 ml RCV media in 20 ml gas chromatography vials (Thermo Fisher Scientific Inc., Waltham, USA). The vials are sealed airtight after inoculation and anaerobised by repeated evacuation of the vial headspace and refill with argon. Cultivation of *R. capsulatus* strains in 96 well microtiter plates (Sarstedt, Nümbrecht, Germany) was conducted in 250 µl RCV in custom build vacuum chambers. The culture media was supplemented to a final concentration of 50 μg/mL kanamycin sulfate (Km), 34 μg/mL chloramphenicol (Cm) or 20 μg/ml streptomycin sulfate (Sm) when appropriate.

Plasmids were introduced into a *R. capsulatus* strains by biparental or triparental conjugation adapted from Katzke *et al.*^47^ In short, the plasmid is used to transform the donor *E. coli* strain (ST18 for biparental, DH5α for triparental mating). The donor strain is cultivated on a LB agar plate. Cell mass of the *E. coli* donor strain and from a freshly grown PY agar plate of the recipient *R. capsulatus* strain is resuspended in 1 mL RCV media, respectively. The *R. capsulatus* strain (250 µl) is mixed with the *E. coli* donor [50 µl; helper DH5α strain carrying the plasmid pRK2013 is additionally added for triparental mating (50 µl)] and plated on top of a Whatman® cellulose acetate membrane filter (Cytiva, Wilmington, United States) on a LB agar plate (no antibiotic). The plate is incubated aerobically at 30 °C for 24 h, the cell mass is suspended in 1 ml RCV media and plated on PY agar plates containing 50 µg/ml Km and 20 µg/ml Sm for selection.

All plasmids used and created in this study are listed in Supplementary Table 2. The primers used in this study were purchased from IDT DNA (Coraville, USA) or Eurofins Genomics (Ebersberg, Germany) and are listed in Supplementary Table 3-6. Polymerase chain reactions (PCRs) were conducted with Q5® High-Fidelity DNA Polymerase (New England Biolabs, Ipswich, USA), PCR purifications with the Monarch® PCR & DNA cleanup kit (New England Biolabs, Ipswich, USA), extraction of genomic DNA with the Monarch® Genomic DNA Purification Kit (New England Biolabs, Ipswich, USA), Gibson assemblies with the NEBuilder® HiFi DNA Assembly Master Mix (New England Biolabs, Ipswich, USA) and Golden Gate cloning with T4 Ligase, BsmBI restriction enzyme or BsaI restriction enzyme (New England Biolabs, Ipswich, USA) according to the instructions provided by the manufacturer. Successful assembly of desired vectors was verified by Sanger sequencing through Microsynth Seqlab GmbH (Göttingen, Germany).

For the generation of plasmid pMM0215, pGGAselect was linearized by PCR using Primers P1 and P2. The *anfD* gene was PCR amplified from pMM0128 with P3 and P4. The purified fragments were cloned via Gibson assembly, and the reaction mix was used to transform chemo competent DH5α cells. The outgrowth was plated on a Luria Broth (LB) agar plate containing 34 µg/ml Cm for selection.

The plasmid pMM0208 was generated by introducing the *lacZα* cassette of pOGG024 into the *anfHDGK* operon in pMM0128. The *lacZα* cassette was PCR amplified with primers P7 and P8 and pMM0128 linearized by PCR using the primers P5 and P6. The amplicons were purified and then by Gibson assembly. The reaction mix was used to transform chemo competent DH5α cells. The outgrowth was plated on LB agar plates containing 50 µg/ml Km, 20 µg/ml X-gal and 100 µM IPTG for selection.

The plasmid pMM0216 was generated by linearizing pMM0215 by PCR using P9 and P10. A mismatch in P9 introduce the α-S(TCG)373S(AGC) mutation to remove a BsmBI cutting site and the primer extensions introduce BsmBI cutting sites with matching 4 bp overhangs. The amplicon was purified and circularized by Golden Gate cloning. The reaction mix was used to transform chemo competent DH5α cells. The outgrowth was plated on LB agar plates containing 34 µg/ml Cm for selection.

The plasmids pMM0128 V1-V3 were isolated from *R. capsulatus* MM0297 V1-V3 obtained from the directed evolution libraries. The plasmids pMM0216 V1-V3 were prepared analogues to pMM0215 using pMM0128 V1-V3 as the *anfD* template. A Strep-tag II was introduced to plasmid pMM0216 V1-V3 by linearizing the plasmids with primers P20 and P21. The amplicons were purified and circularized by Golden Gate cloning. The reaction mix was used to transform chemo competent DH5α cells. The outgrowth was plated on LB agar plates containing 34 µg/ml Cm for selection. pMM0216 V1-V3 Strep-tag II were subcloned into pMM0208 to generate pMM0128 V1-V3 Strep-tag II by Golden Gate cloning.

For the generation of pMM0128 V57G*^anfD^*, pMM0128 α-V57A*^anfD^*, pMM0128 α-H180Q*^anfD^* and pMM0128 V57A*^anfD^*/H180Q*^anfD^* and pMM0128 V57G*^anfD^*/H180Q*^anfD^* plasmid pMM0128 was amplified by PCR using the primers P11-P15. The primer extensions introduce BsaI cutting sites and matching 4 bp overhangs. The amplicon was purified and ligated by Golden Gate cloning. The reaction mix was used to transform chemo competent DH5α cells. The outgrowth was plated on LB agar plates containing 50 µg/ml Km for selection. Plasmid pMM0132 V82A*^nifD^*, pMM0132 H207Q*^nifD^*and pMM0132 V82A*^nifD^*/H207Q*^nifD^* were prepared analogously using pMM0132 as template and primers P16-P19. The

#### 1.1.3 Library generation and directed evolution cycle

Initial library generation of *anfD* in pMM0128 was performed using megaprimer PCR with primers P22 and P95.^48^ This approach was later adapted to a two-stage Golden Gate cloning–based method to improve robustness. The site-saturation mutagenesis at targeted site of *anfD* were introduced in plasmid pMM0216 by PCR using primers with extensions that contain the BsmBI cutting site, matching 4 bp overhangs and the NNK degenerate codon at the desired position (P96-P163). The amplicon was purified and ligated using Golden Gate cloning. The Golden Gate reaction mix (final volume 20 μL) contained T4 ligase buffer (2 μL), T4 ligase (0.5 μL), BsmBI (0.5 μL) and the PCR amplicon (150 ng) in ddH_2_O. The mix was incubated in a thermocycler: 30x (5 min at 42 °C; 5 min at 16°C), 5 min at 60 °C, stored at 12 °C. The reaction mix was used to transform chemo competent DH5α cells. The outgrowth was plated on LB agar plates containing 34 µg/ml Cm for selection. The resulting single colonies were pooled and the plasmid is isolated. After the mutagenesis of the target site was confirmed by Sanger sequencing, the plasmid pMM0216 mutant library was cloned into pMM0208 to generate the mutant library of pMM0128 by Golden Gate cloning using the BsaI restriction sites. The Golden Gate reaction mix (final volume 20 μL) contained T4 ligase buffer (2 μL), T4 ligase (0.5 μL), BsaI (0.5 μL), pMM0208 (100 ng) and the pMM0216 library (100 ng) in ddH_2_O. The mix was incubated in a thermocycler: 30x (5 min at 37 °C; 5 min at 16°C), 5 min at 60 °C, stored at 12 °C. The reaction mix was used to transform chemo competent DH5α cells. The outgrowth was plated on LB agar plates containing 50 µg/ml Km, 20 µg/ml X-gal and 100 µM IPTG for selection. The single colonies were pooled and suspended 1 ml LB media and used to prepare a glycerol stock (adding 20% V/V glycerol).

The *E. coli* DH5α carrying the pooled pMM0128 library was used for conjugation into *R. capsulatus* expression strain MM0167 by triparental mating. The conjugation cell mixture is selected on PY agar plates supplemented with 5 PY agar plates containing 50 µg/mL Km and 20 µg/mL Sm under phototrophic growth conditions. Single colonies of MM0164 carrying individual mutant plasmids of the pMM0128 library were inoculated into RCV medium (250 µl, supplemented with 10 mM serine and 50 µg/mL Km) in 96 well microtiter plates using a colony picking robot (PIXL, Singer Instruments, Watchet, England). The plates were transferred into custom build vacuum chambers and placed on top of a water reservoir (48 well plate containing 12 ml of diH_2_O). The atmosphere in the chamber was set to 1.2 atm argon and the plates were cultivated phototrophically for 6 d.

#### 1.1.4 *In vivo* CH_4_ formation assay

The precultures from the 96 well plates were inoculated into GC vials (200 µL inoculated in 4 ml RCV, OD_660_ ∼0.1), the vials were sealed with butyl rubber stoppers and the headspace was exchanged to 8% CO_2_ in argon to a final pressure of 1.2 atm by three evacuation and refill cycles with argon followed by a final cycle with 8% CO_2_ in argon (ARCL™ speed, Air liquide). The assay cultures were incubated for 4 d under phototrophic conditions (LED panels). The assay was stopped by removing the cultures from the light source.

#### 1.1.5 Protein production and purification

The Fe-nitrogenase WT and variants were purified by affinity chromatography. The Fe-nitrogenase was purified according to established protocols.^14^ For the Fe-nitrogenase variants, the *R. capsulatus* expression strains [MM0425 carrying pMM0128 V1-Strep-tag II (MM0598), V2-Strep-tag II (MM0845), V3-Strep-tag II (MM0846)] were inoculated on PY agar plates containing 50 µg/mL Km. The plates were incubated phototrophically for 48 h at 30 °C under an Ar atmosphere. Obtained cell mass was inoculated in argon purged modified RCV medium (50 mL) containing 1 mM Fe(III) citrate as the Fe source. The cultures were cultivated phototrophically at 30 °C for 24 h. Next, the liquid cultures were used to inoculate 1 L of modified RCV medium purged with Ar to an *OD*_660_ of 0.1 for protein production. Protein purification was initiated when the cultures reached an OD_660_ of ∼ 2.5. Isolation of the Fe-nitrogenase variants was performed analogously to the purification procedure that we established previously for the Fe-nitrogenase.^14^ For SDS–PAGE analysis, protein samples were mixed with Pierce™ Lane Marker Reducing Sample Buffer (Thermo Fisher Scientific) and heat-treated at 98 °C for 10 min to induce denaturation. Insoluble material was removed by centrifugation at 16,000 × g for 1 min, after which the clarified samples were applied to 4–20% Mini-PROTEAN TGX Stain-Free gels (Bio-Rad Laboratories, Inc.). PageRuler™ Plus Prestained Protein Ladder (Thermo Fisher Scientific) was used for molecular mass estimation. Electrophoretic separation was carried out at 180 V for 30 min, and protein bands were detected using GelCode™ Blue Safe Protein Stain (Thermo Fisher Scientific).

#### 1.1.6 *In vitro* nitrogenase assays

The specific nitrogenase activities were measured for the formation of H_2_, NH_3_, CO, CH_4_, HCOO^-^, C_2_H_4_ and C_2_H_6_ under 1.2 atm of N_2_, CO_2_ or 10% C_2_H_2_ in Ar. Assays were set up under an Ar atmosphere by adding nitrogenase catalytic component (0.1 mg) to an anaerobic solution of 50 mM TRIS (pH = 7.8), 10 mM sodium dithionite, 3.5 mM adenosine triphosphate (ATP), 7.87 mM MgCl_2_, 44.59 mM creatine phosphate and 0.20 mg/mL creatine phosphokinase (catalogue number: C3755; Sigma-Aldrich St. Louis, USA). The reaction vials were sealed with butyl rubber stoppers, and the headspace was exchanged to 1.2 atm N_2_, Ar or CO_2_. Following a 10 min incubation at 30 °C, the reactions were initiated by adding Fe-nitrogenase reductase component (0.5 mg) to a final volume of 700 µL. Reactions were incubated at 30 °C and moderate shaking (300 rpm) for 9 min before quenching with 300 µL of 400 mM sodium ethylenediaminetetraacetic acid solution (pH = 8.0).

#### 1.1.7 Diazotrophic growth curves

*R. capsulatus* Fe-nitrogenase expression strains (MM0468, MM0598, MM0845, MM0846) were inoculated from freshly grown PY-agar plates into 96-well microtiter plates (250 µL RCV medium supplemented with 3.5 mM serine). After 6 d cultivation under anaerobic and phototrophic conditions, the precultures were inoculated into GC vials (4 mL N-free RCV medium) and sealed airtight with butyl rubber stoppers. The medium and headspace of the culture flasks were purged with N_2_ and set to a final pressure of 1.2 atm. The bacteria were cultivated phototrophically at 30 °C. Samples for OD_660_ measurements were taken at the indicated time points by piercing the septum of the cultivation vials with a syringe. The OD_660_ was measured with an Infinite® 200 PRO plate reader (Tecan Group Ltd, Männedorf, Switzerland).

#### 1.1.8 Quantification of CH_4_, H_2_, CO. C_2_H_4_ and C_2_H_6_

The amount of CH_4_ formed in the *in vivo* CH_4_ assay was determined through headspace analysis using a PerkinElmer® Clarus®690 GC system (GC–FID/TCD) with a custom-made column circuit (ARNL6743). The headspace samples are injected by a TurboMatrix X110 (PerkinElmer Inc., Waltham, USA) auto sampler heating the samples to 45 °C for 15 minutes prior to the injection. The samples were then separated on a RGAP column (7’ RGAP 1/8“ Sf; PerkinElmer Inc., Waltham, USA) followed by a RGA20 (9 17’ RGA20, 1/8” Sf; PerkinElmer Inc., Waltham, USA), RGAP (9 17’ RGA20, 1/8“ Sf; PerkinElmer Inc., Waltham, USA), and RGA column (7” RGA, 1/8“ Sf; PerkinElmer Inc., Waltham, USA) all kept at 60 °C. Subsequently, the gases were passed again over the RGAP column before running over an ASA column (4’ ASA, 1/8” Sf, PerkinElmer Inc., Waltham, USA). The gases are then detected with a flame ionization detector (FID, at 250 °C, time per sample: 4 min). The quantification of CH_4_ was based on a linear standard curve that was derived from measuring varying amounts of CH_4_ using the same instrument.

Simultaneous quantification of H_2_, CH_4_, C_2_H_4_, C_2_H_6_ and CO was conducted with the same instrument. The samples were separated on a HayeSep column (7′ HayeSep N 1/8″ Sf; PerkinElmer Inc., Waltham, USA), followed by a molecular sieve (9′ Molecular Sieve 13× 1/8″ Sf; PerkinElmer Inc., Waltham, USA) kept at 60°C. Subsequently, the gases were detected with an FID (at 250°C) and a TCD (at 200°C). The quantification of all substrates was based on linear standard curves derived from measuring varying amounts of CO, CH_4_, H_2_, C_2_H_4_ and C_2_H_6_ under identical conditions (time per sample: 15 min). The results were plotted using the GraphPad Prism 9 software (Dotmatics, Boston, USA).

#### 1.1.9 Quantification of NH_3_

NH_3_ was quantified with fluorescence-based assay as described by Corbin.^49^ 10 µL sample were combined with 90 µL of a solution containing 2 mM *o*-phthalaldehyde, 10 % (V/V) ethanol, 0.05 % (V/V) β-mercaptoethanol and 0.18 M potassium phosphate buffer (pH = 7.3) and incubated at 25 °C for 2 h in the dark in Nunc™ F96 MicroWell™ plate (Thermo Fisher Scientific Inc., Waltham, USA). Fluorescence at 485 nm was monitored with an Infinite® 200 PRO plate reader (Tecan Group Ltd, Männedorf, Switzerland) in fluorescence top reading mode using an excitation wavelength of 405 nm. The quantification of NH_3_ was based on a linear standard curve that was derived from measuring varying amounts of NH_4_Cl under identical conditions. Samples incubated under a CO_2_ atmosphere instead of dinitrogen were used to correct for background signal. The results were plotted using the GraphPad Prism 9 software (Dotmatics, Boston, USA).

#### 1.1.10 Quantification of formate via GC-MS

Formate quantification via gas chromatography-mass spectrometry (GC-MS) was performed according to ^50^. For derivatization, 0.1 mL of sample was combined with 0.5 mL of a 100 mM pentafluorobenzyl bromide solution and 0.1 mL of 500 mM potassium phosphate buffer (pH 6.8). The mixture was thoroughly vortexed and incubated at 60 °C for 1 h with agitation at 500 rpm. After cooling to ambient temperature, 1 mL of a 100 µM solution of 1,3,5-tribromobenzene in *n*-hexane was added. The samples were mixed again, followed by phase separation via centrifugation at 1,400 × g for 15 min. An aliquot (300 µL) of the organic layer was transferred to a 1.5 mL short-thread vial (VWR, Radnor, USA).

The formate content was analysed on an Agilent 8860 gas chromatograph (split = 1:20), equipped with an HP5 MS column that was coated with dimethylpolysiloxane (30 m, 0.25 mm i.d., 0.25 µm film thickness; Agilent, Santa Clara, CA, USA). The gas chromatograph was connected to an Agilent 5977 mass spectrometer (Agilent, Santa Clara, CA, USA) and the system was operated with the Mass Hunter Workstation GC/MS Data Acquisition Software ver. 10.2 (Agilent, Santa Carla, CA, USA). The column’s initial temperature (50 °C) was held for 3 min and then increased to 280 °C at a rate of 20 °C/min. The injection port was kept at 280 °C and ion source at 230 °C. Helium was used as the carrier gas at a flow rate of 1.5 mL/min. Mass spectra were obtained by positive-ion electron ionization (EI) mode scanning every 0.74 s from 40 – 600 *m/z*. Selected ion recording (SIR) was measured every 0.12 s for the molecular peak ion of the derivative of HCOO^-^ at 226 *m/z* and the base peak ion of 1,3,5-tribromobenezene at 314 *m/z.* The ionization energy of the EI condition was 70 eV. A linear standard curve (R^2^ ≥ 0.998) for HCOO^-^ was generated by plotting the ratio of the peak at 226 *m/z* and 314 *m/z* against the concentration of HCOO^-^. The results were plotted using the GraphPad Prism 9 software (Dotmatics, Boston, USA).

#### 1.1.11 Cryo-EM sample preparation, data collection and processing

The trapping of the Fe-nitrogenase V3 complex Anf(DGK)_2_(H_2_)_2_ was conducted as described in ref. ^51^. Four µL of the V3 complex (0.5 mg ml^−1^) were applied to glow-discharged QUANTIFOIL R2/1 300 copper mesh grids (Quantifoil Micro Tools) and blotted for 10 s with a blot force of 4 at ∼90% humidity and 4 °C using a Vitrobot Mark IV (Thermo Fisher Scientific, Waltham, USA) that was placed inside an anaerobic COY tent. Cryo-EM grids were vitrified by plunge freezing into a liquid ethane/propane mixture (37/63, v/v) and stored in liquid nitrogen prior to imaging. Data acquisition was performed on a Titan Krios G3i electron microscope (Thermo Fisher Scientific, Waltham, USA) operated at an acceleration voltage of 300 kV and equipped with a BioQuantum K3 energy filter (Gatan, Pleasanton, USA). Images were recorded in electron-counting mode at a nominal magnification of 105,000×, corresponding to a pixel size of 0.837 Å, with a total electron exposure of 50 e^−^/Å^2^ fractionated (50 fractions). Aberration-free image shift correction was applied during data collection using EPU software (versions 2.9–2.11; Thermo Fisher Scientific, Waltham, USA), and defocus values ranged from −1.4 µm to −2.4 μm.

The data set was processed in cryoSPARC v.4.1(Extended Data Fig. 7,8).^52^ Dose-fractionated movies were gain-normalized, aligned and dose-weighted by Patch Motion Correction. The contrast transfer function (CTF) was calculated by the Patch CTF routine. Next, 1,592,134 initial particles were extracted using the blob picker with a box size of 448 pixels, which were downsampled to a box size of 192 pixels. These were used to build two-dimensional (2D) classes. After selection of 21 2D classes, ab initio reconstruction was performed with 100,000 particles generating four three-dimensional (3D) classes. Three of the 3D classes were used for a heterogeneous refinement of the 1,592,134 initial particles. The largest 3D class contained 923,185 particles exhibited nitrogenase like features and was used to create templates for template picking. The template picking extracted 1,582,288 particles at a box size of 448 pixels, which were used for 2D classification. 34 of 50 2D classes were selected and heterogeneous refinement was performed using the previous volumes generating a 3D class with the expected topology (756,049 particles). For further cleaning, 100,000 particles were used for ab initio reconstruction and classification in three 3D classes. Two of their volumes were used to perform heterogeneous refinement generating a nitrogenase like 3D class (716,595 particles). The class was used for non-uniform refinement with C2 symmetry, yielding a density map with a 2.52-Å gold-standard Fourier shell correlation (GSFSC) split resolution. The volume was used for reference motion correction and the resulting class (714,366 particles was subjected to local CTF refinement, per-particle defocus optimization, and Ewald sphere (EWS) correction yielding a density map with a GSFSC resolution of 2.04 Å.

#### 1.1.12 Model building and refinement

The initial cryo-EM map fitting was conducted using the structure of the WT Fe-nitrogenase Anf(DGK)_2_(H_2_)_2_ complex (pdb: 8OIE) in Chimera 1.7.1.^53^ Automatic refinement of the structure was done using phenix.real_space_refine of the Phenix v1.21.1 software (Extended Data Fig. 8).^54^ Manual refinements and water picking were performed with Coot v0.8.9.2.^55^

#### 1.1.13 Protein mass spectrometry and peptide identification

For mass spectrometric analysis, a solution of 30 µg purified Anf(DGK)_2_ (V3) in 1.3% deoxycholic acid (DOC) and ammonium bicarbonate buffer was carbamidomethylated with TCEP and iodacetamide. After diluting the DOC to a final concentration of 0.3%, 0,6 µg trypsin (SERVA, sequencing grade, modified) was added and incubated for 1 hour at 30°C. The sample was then acidified with trifluoroacetic acid (TFA) and desalted using C18-SPE, dried in a Speedvac and reconstituted in 0.1% TFA.

The peptides were analysed using liquid chromatography-mass spectrometry in an Orbitrap Exploris 480 (Thermo Fisher Scientific) coupled to a Thermo Scientific Vanquish UHPLC system. For an LCMS run, 0.2 µg of peptides were injected and separated on a C18 reversed-phase HPLC column using a 30-minute gradient (0.15% formic acid to 0.15% formic acid with 35% acetonitrile). The data were recorded in DDA mode and analysed using Proteome Discoverer software (Thermo Scientific) and Byonic software (Protein Metrics) and the sequence data files from *R. capsulatus* SB1003 (and the V3 sequence).

Only trypsin/P specificity, up to two missing cleavages, fixed carbamidomethylation and variable methionine oxidation were considered. The precursor mass tolerance was 10 ppm. A Byonic Wildcard search was added for methionine in the range of-30 to +100 Da to identify potential modifications at residue M362*^anfD^*.

## Acknowledgments

The authors thank the Rebelein laboratory for fruitful discussions and valuable comments on the manuscript. We thank B. Masepohl and T. Drepper for providing strains and plasmids.

## Author contributions

J.G.R. conceived and supervised the project. J.G.R. acquired funding. N.N.O. and J.G.R. designed and analysed experiments. N.N.O. performed molecular work. N.N.O., J. C. and F.V.S. performed anaerobic protein purification. N.N.O. performed *in vitro* enzyme biochemistry. N.N.O and P.C. performed product quantification. N.N.O. performed *in vivo* experiments. N.N.O. J.C. and J.K. performed mass spectrometry analysis. F.V.S., N.N.O and S.P. performed cryo-EM data acquisition. N.N.O and J.Z. processed and refined the cryo-EM structure with support of T.J.E. N.N.O, J.Z. and J.G.R. analysed the cryo-EM structure. N.N.O. and J.G.R. wrote the original manuscript, which was reviewed and edited by all authors.

## Data and materials availability

All unique materials used in this study are available from the corresponding author upon request. All raw data for *in vivo* work, kinetic experiments and protein characterizations will be deposited on Edmond, the Open Research Data Repository of the Max Planck Society.

---

**Extended data Fig. 1.**
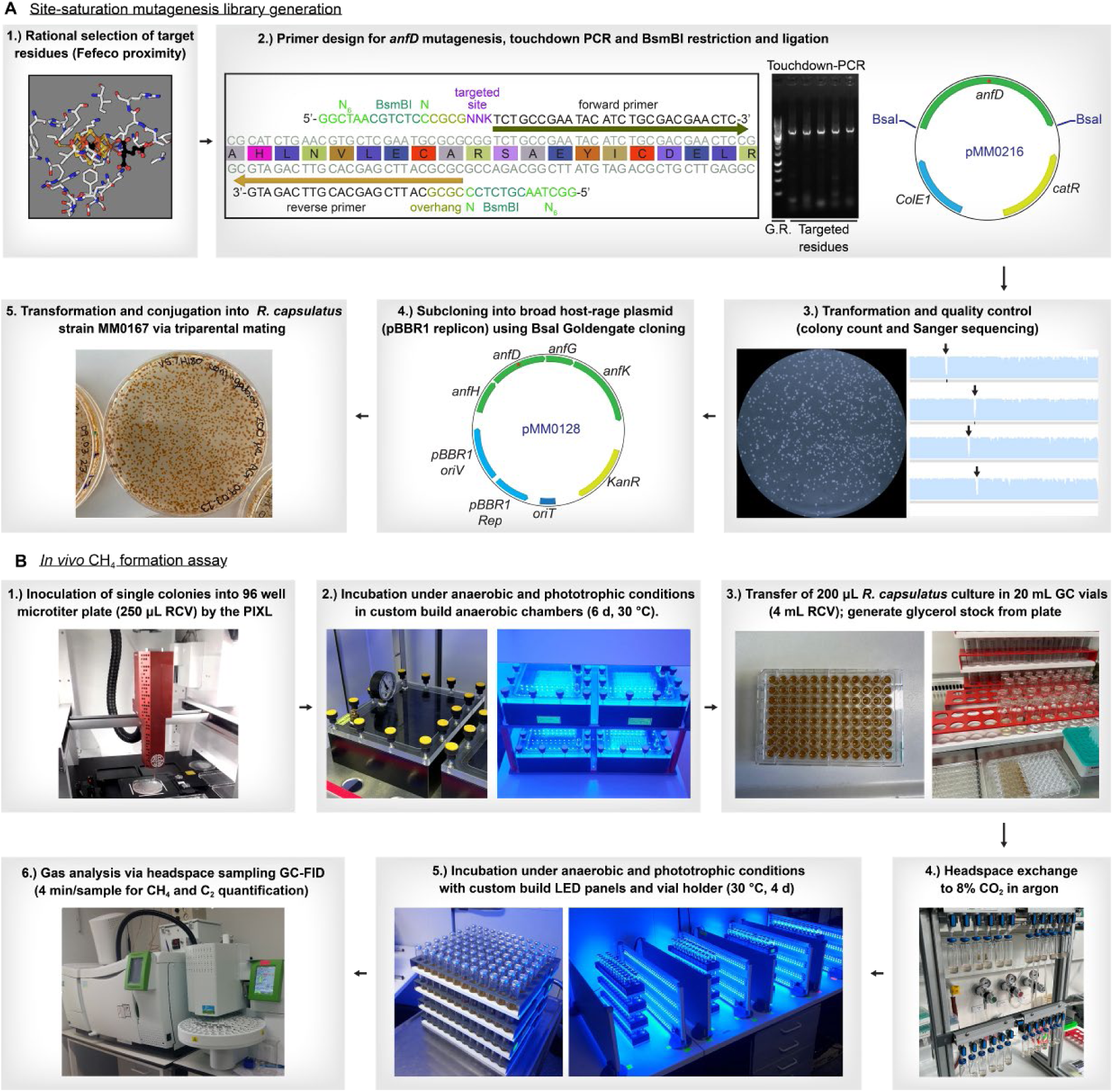
Directed evolution workflow. (A) Site-saturation mutagenesis library generation. Sites in *anfD* are selected based on their proximity to the active site cofactor FeFeco using the structure determined by cryo–electron microscopy (PDB: 8OIE).^51^ Mutagenesis primers are designed to introduce a NNK degenerate codon and the BsmBI restriction site via primer overhangs. The site-saturation library is generated by amplifying the *anfD* template plasmid (pGGAselect *anfD* cloning vector, silent mutation α-S(TCG)373S(AGC) removes BsmBI cutting site, Cm^R^, *ColE1* replicon) by touchdown PCR. The PCR product formation is confirmed by agarose gel electrophoresis (G.R.: GeneRuler 1 kb DNA Ladder, Thermo Fisher Scientific). The purified DNA fragment is circularized by restriction and ligation cloning in a Golden Gate–like cloning reaction to form the mutated plasmid pMM0216. The reaction is used to transform *E. coli* DH5α and the library quality is assessed by colony counts and Sanger sequencing of the pooled library. Library formation is indicated by a decrease in coverage (black arrows). pMM0216 is subcloned into the broad-host-range plasmid pMM0208 by Golden Gate cloning using the BsaI restriction enzyme to form the Fe-nitrogenase expression plasmid pMM0128 (pBBR1 replicon). The golden gate reaction is used to transform *E. coli* DH5α, the library is pooled and conjugated into *R. capsulatus* strain MM0167 by triparental mating. (B) CH_4_ assay. Single colonies of *R. capsulatus* are inoculated into a 96-well microtiter plate (250 µL RCV) using a colony-picking robot (PIXL). The plate is transferred into a custom-built vacuum chamber with a transparent lid [poly(methyl methacrylate)] on top of a water reservoir plate (48 well plate containing 12 mL water). The atmosphere of the chamber is exchanged for 8% CO_2_ in Ar and the plates are incubated for 6 d at 30 °C illuminated by custom-built LED panels (λ: 850 nm, 420 nm and 470 nm). The preculture plate is used to inoculate 200 µL culture into 3.8 mL RCV in GC vials. Glycerol (75 µL, 80% v/v in H_2_O) is added to the remaining *R. capsulatus* cultures for a cryogenic glycerol stock. The GC vials are capped airtight with butyl rubber stoppers and the headspace is exchanged to 8% CO_2_ in Ar. The CH4 assay cultures are incubated in custom-built vial holders at 30 °C illuminated by LED panels (λ: 850 nm, 420 nm and 470 nm, ∼60 μmol photons m^-2^ s^-1^). The final yield of CH_4_ and C_2_H_6_ is quantified by GC-FID after 4 d of incubation.

**Extended data Fig. 2.**
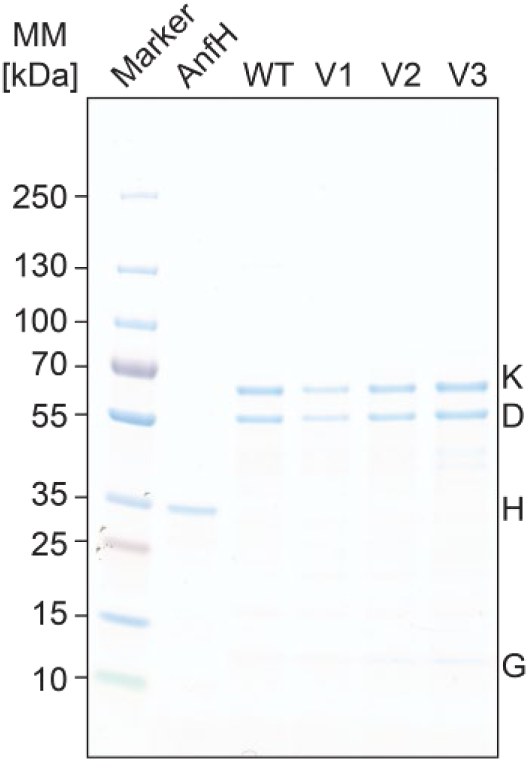
SDS-PAGE analysis. SDS-PAGE analysis of the purified Fe-nitrogenase catalytic component [Anf(DGK)_2_] and the catalytic component of the mutants V1-3.

**Extended data Fig. 3.**
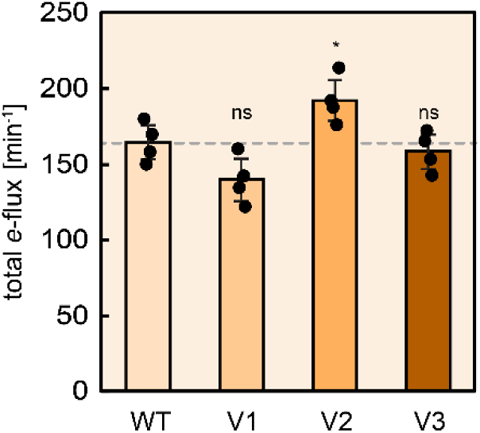
Total electron flux of the Fe-nitrogenase variants. Sum of electron flux into the reaction products of the Fe-nitrogenase WT and mutants in the in vitro assays of Fig. 3 (H_2_: 2 e^−^, CO: 2 e^−^, HCOO^−^: 2 e^−^, CH_4_: 8 e^−^). Each dot represents an activity assay. n = 4 independent experiments. Statistical analysis was performed using unpaired two-tailed Welch’s t-tests against the WT: ns: not significant, *P ≤ 0.05.

**Extended Data Fig. 4.**
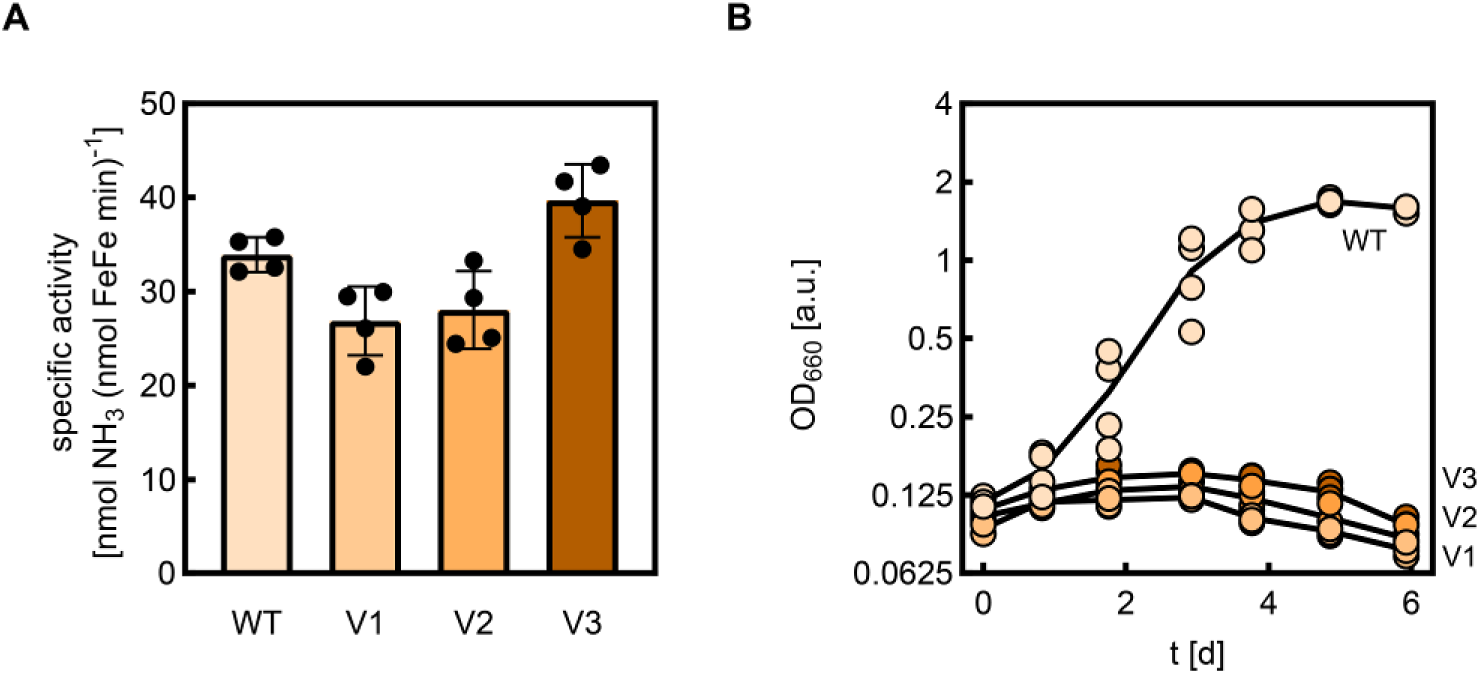
Nitrogen fixation by the Fe-nitrogenase variant. (A) Product formation rates for *in vitro* activity assays conducted under a pure N_2_ atmosphere (1.2 atm) using 0.1 mg Anf(DGK)_2_ and 0.5 mg (20× molar excess) AnfH_2_. Plotted are specific activities for the formation of NH_3_. Each dot represents an activity assay. n = 4 independent experiments. (B) Diazotrophic growth curves of *R. capsulatus* strains expressing the Fe-nitrogenase variants. Each dot represents the values from biological replicates (*n* = 4).

**Extended Data Fig. 5.**
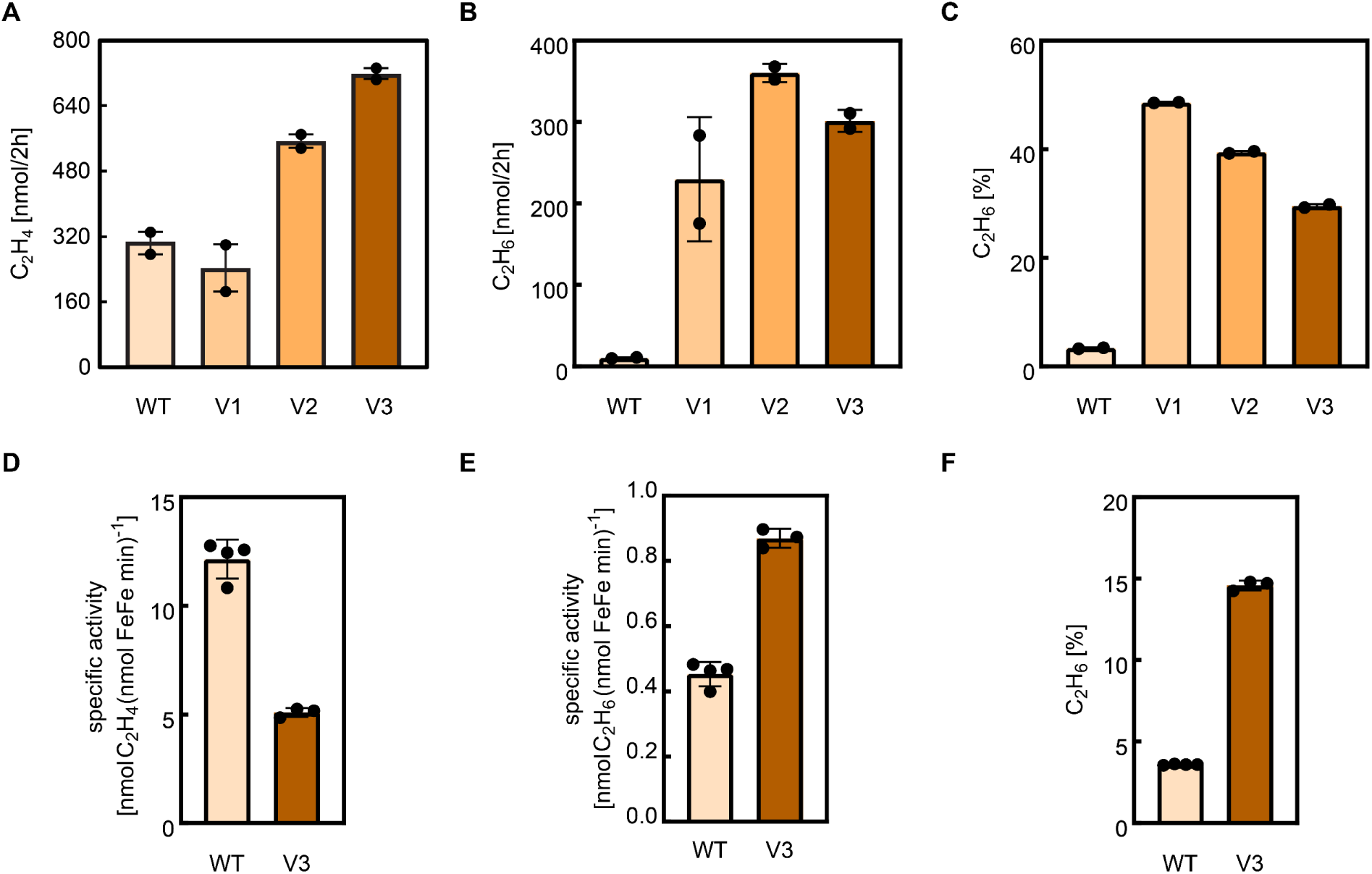
Acetylene reduction by the Fe-nitrogenase variants. C_2_H_4_ (A) and C_2_H_4_ (B) formation in *in vivo* C_2_H_2_ reduction assays conducted under a 1.2 atm 10% C_2_H_2_ in Ar atmosphere with 2 mL *R. capsulatus* culture expressing the Fe-nitrogenase variants. Plotted are the final yield of C_2_H_4_ and C_2_H_6_ per culture after 2 h of incubation. Each dot represents an activity assay. n = 2 independent experiments. (C) Percentage of C_2_H_6_ of the C_2_H_2_ reduction products formed *in vivo* (C_2_H_4_ and C_2_H_6_). (D-F) Product formation rates for *in vitro* activity assays conducted under a 10% C_2_H_2_ in Ar atmosphere using 0.1 mg Anf(DGK)_2_ and 0.5 mg (20× molar excess) AnfH_2_. Plotted are specific activities for the formation of C_2_H_4_ (D) and C_2_H_6_ (E). Each dot represents an activity assay. n = ≥3 independent experiments. (F) Percentage of C_2_H_6_ of the C_2_H_2_ reduction products formed *in vitro* (C_2_H_4_ and C_2_H_6_).

**Extended Data Fig. 6.**
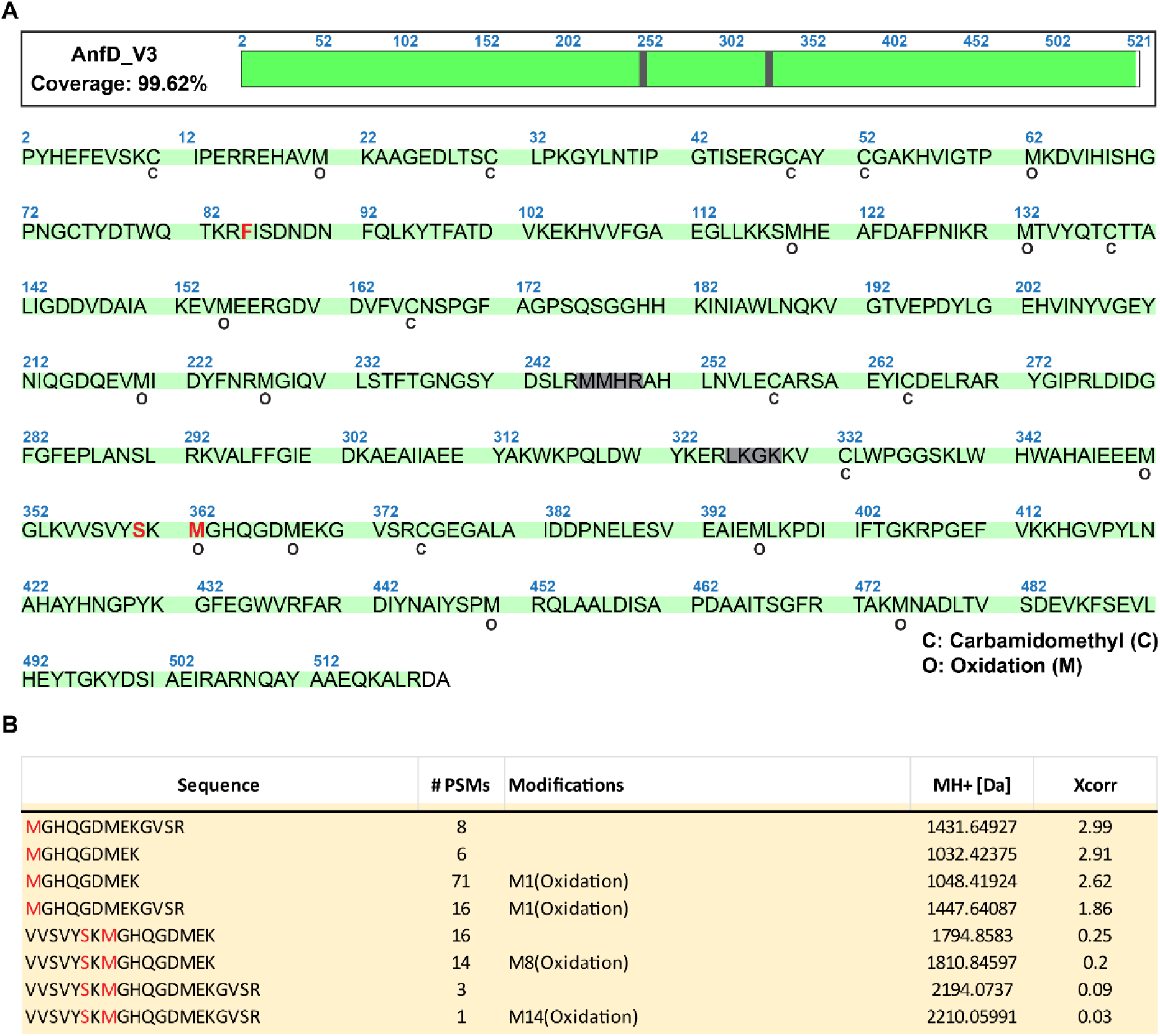
Peptide identification for V3 by mass spectrometry. (A) Coverage of AnfD*^V3^* in the peptide identification. The mutations of V3 are highlighted in red. The identified residue modifications are indicated by **o** (oxidation) and **c** (carbamidomethylation). (B) List of identified peptides including the residue M362*^V3^*. Oxidation of the methionine was the only modification found for M362.

**Extended Data Fig. 7.**
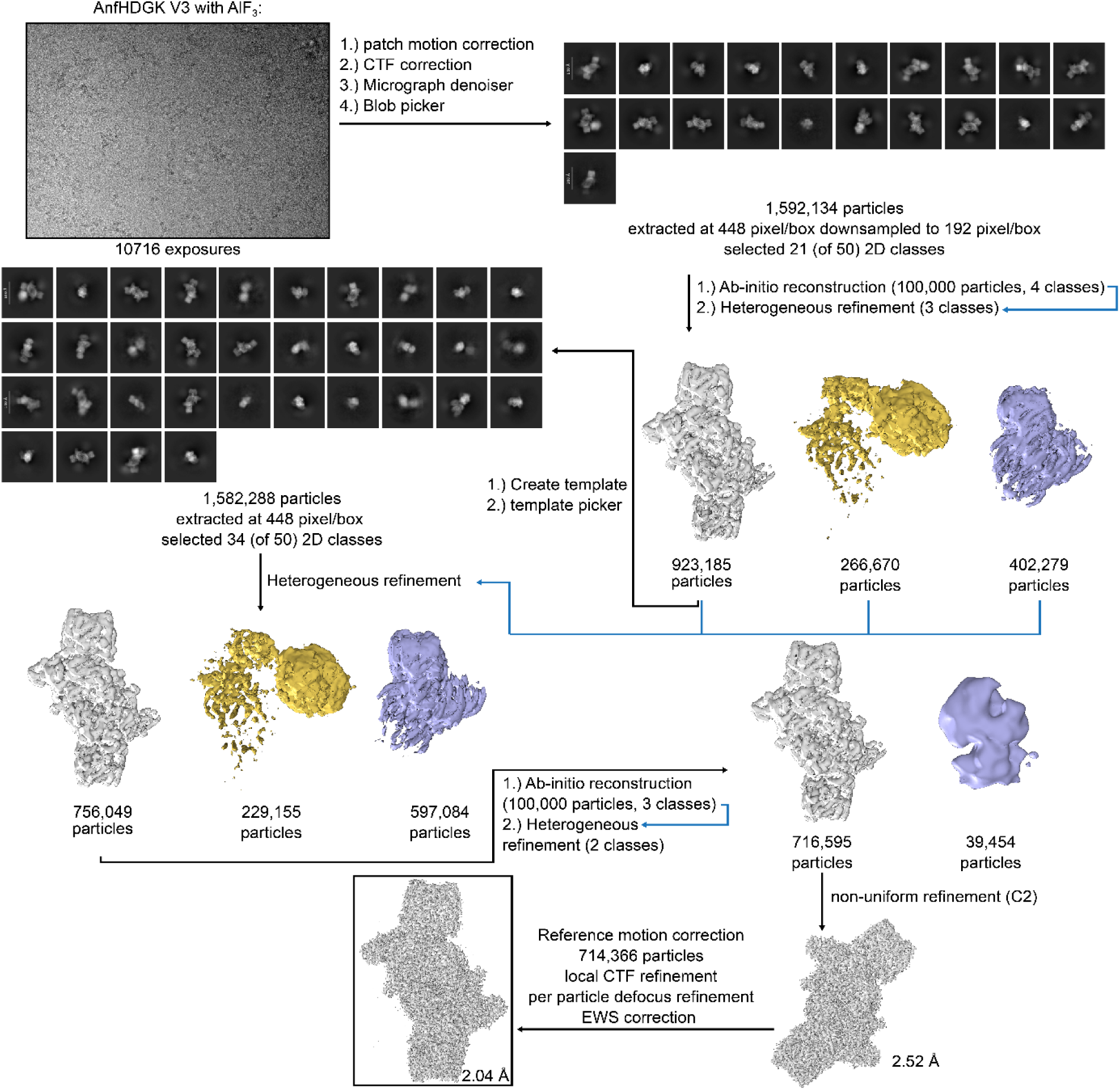
Cryogenic electron microscopy data collection and processing of V3 Anf(DGK)_2_(H_2_)_2_. Schematic overview of the data-processing workflow used to generate the electron density map of V3. Data were collected on a Titan Krios G3i electron microscope operated at 300 kV and equipped with a BioQuantum energy filter and a K3 direct electron detector. All processing steps were performed in CryoSPARC.^52^

**Extended Data Fig. 8.**
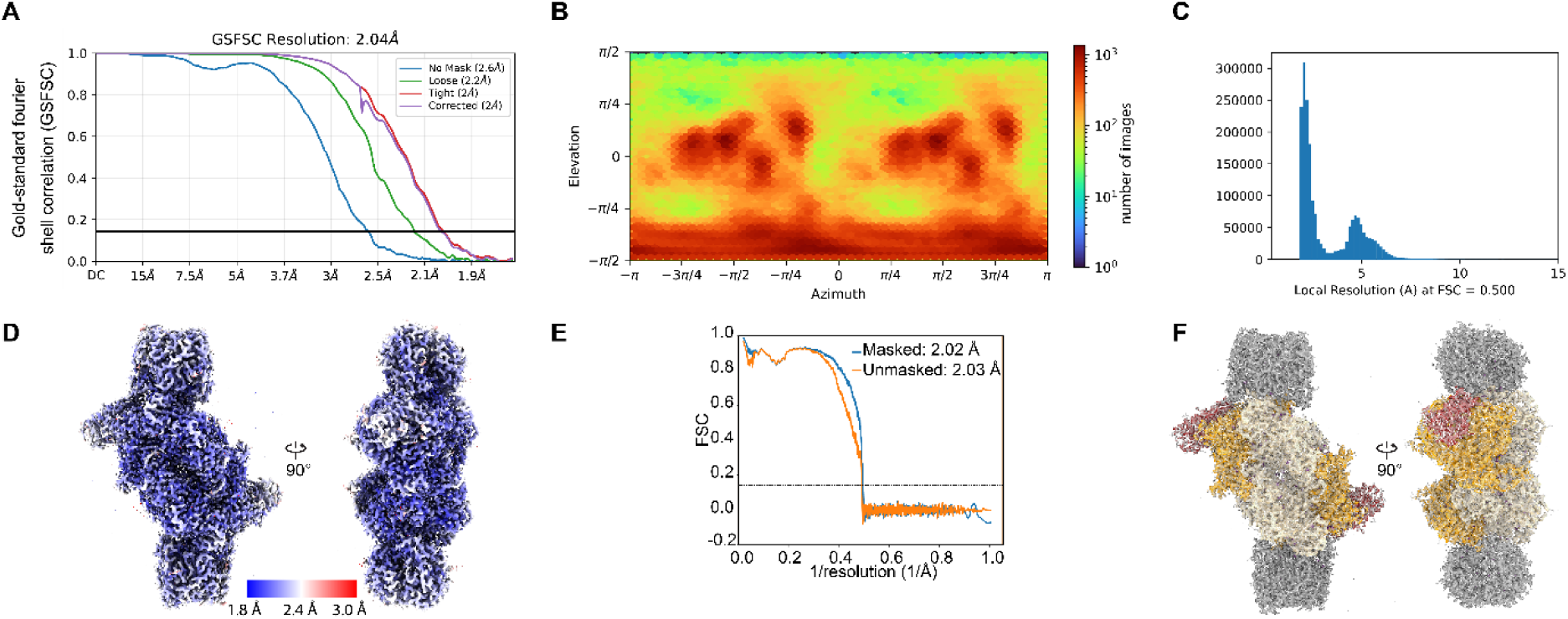
Cryogenic electron microscopy data analysis and model building of V3 Anf(DGK)_2_(H_2_)_2_. (A) Gold-standard Fourier shell correlation curve obtained during map refinement in CryoSPARC, with the resolution estimated at an FSC cutoff of 0.143. (B) Angular distribution of particle orientations. (C) Local resolution distribution calculated using an FSC threshold of 0.143. (D) CryoSPARC-derived local resolution mapped onto the refined density, shown from multiple viewpoints. (E) Fourier shell correlation between the refined map and atomic model, using an FSC cutoff of 0.5. (F) Model-to-map fit displayed from two orientations.

**Table S1.**
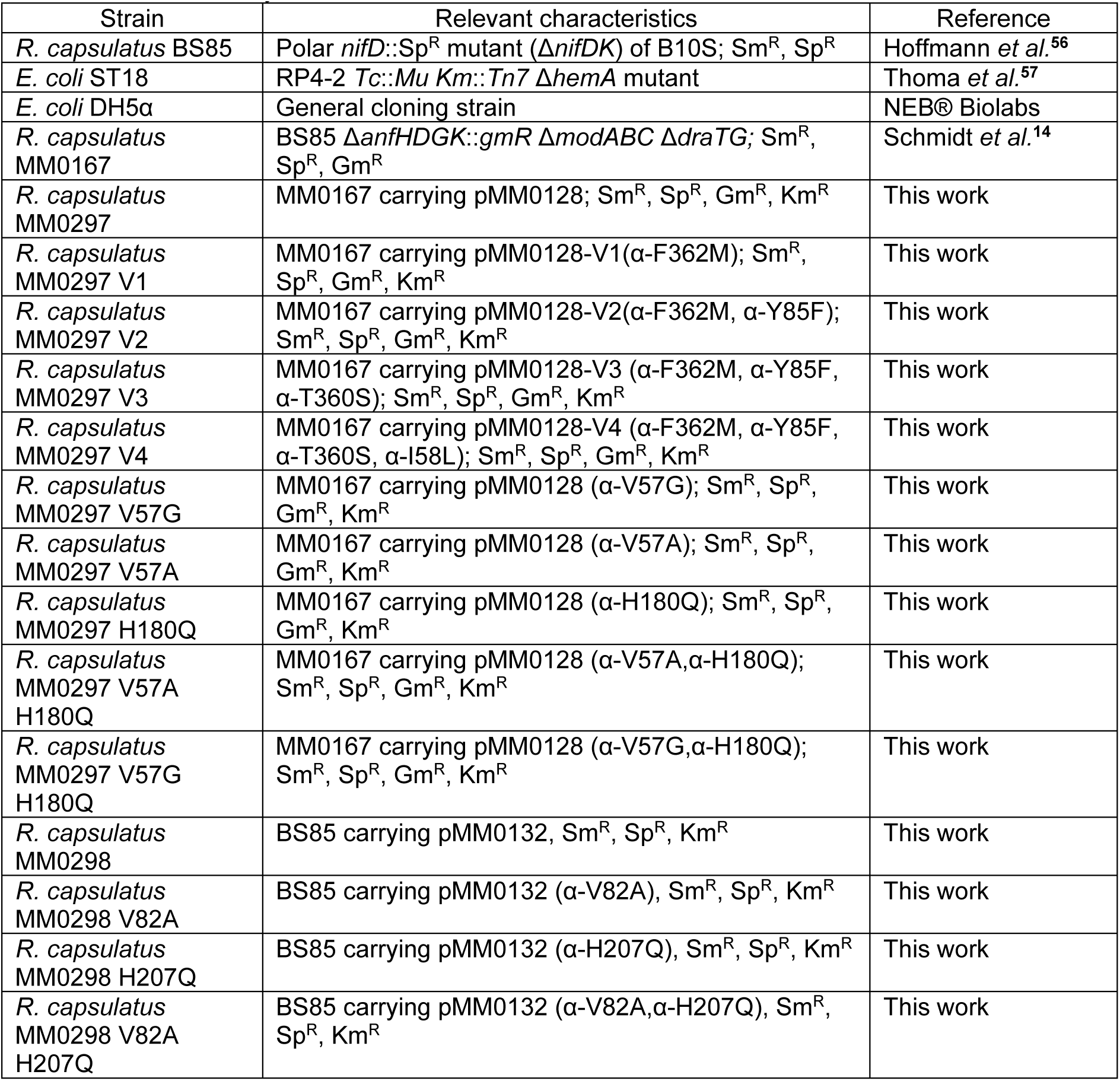
Strains used in this study.

**Table S2.**
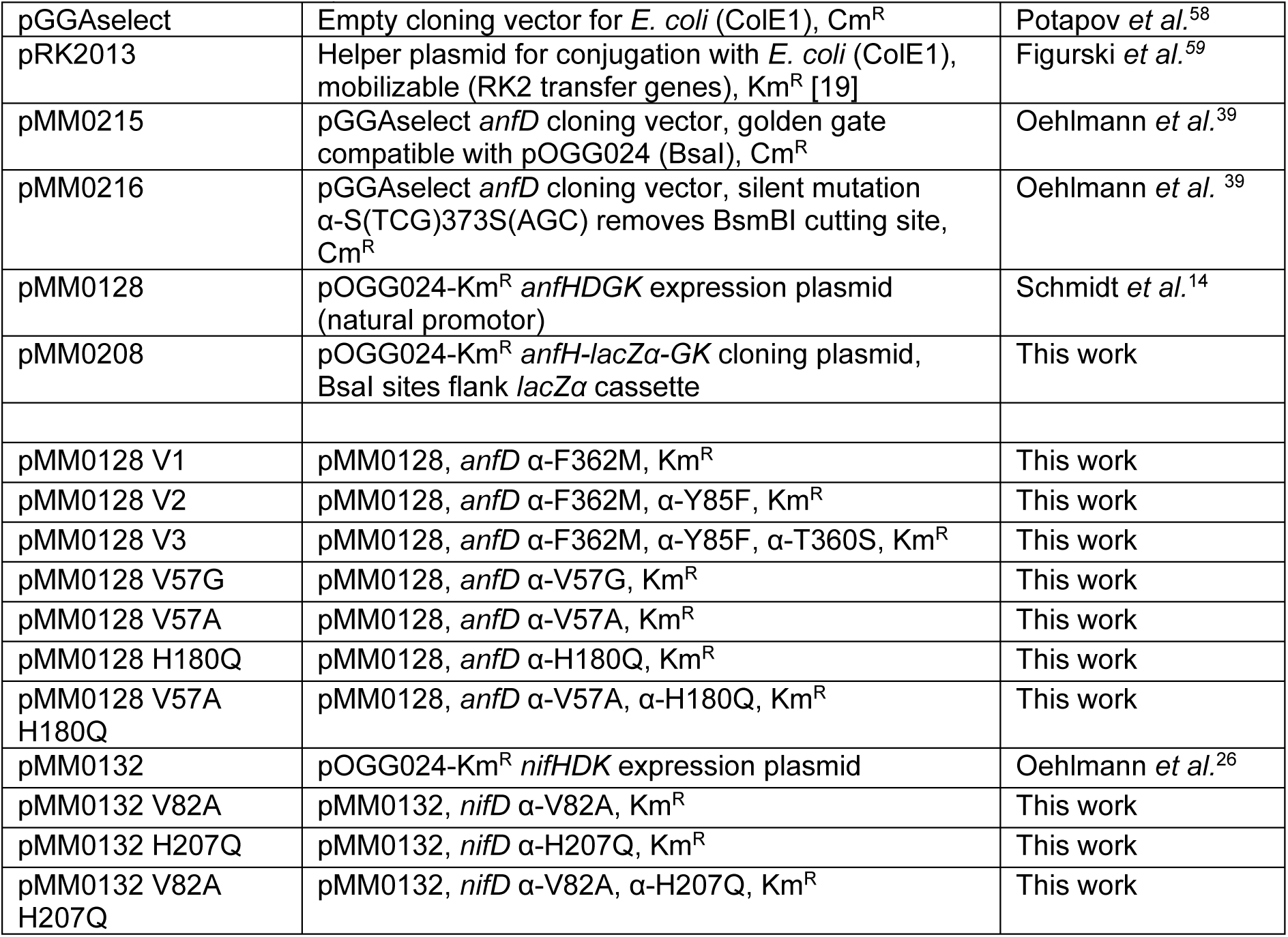
Plasmids used in this study.

**Table S3.**
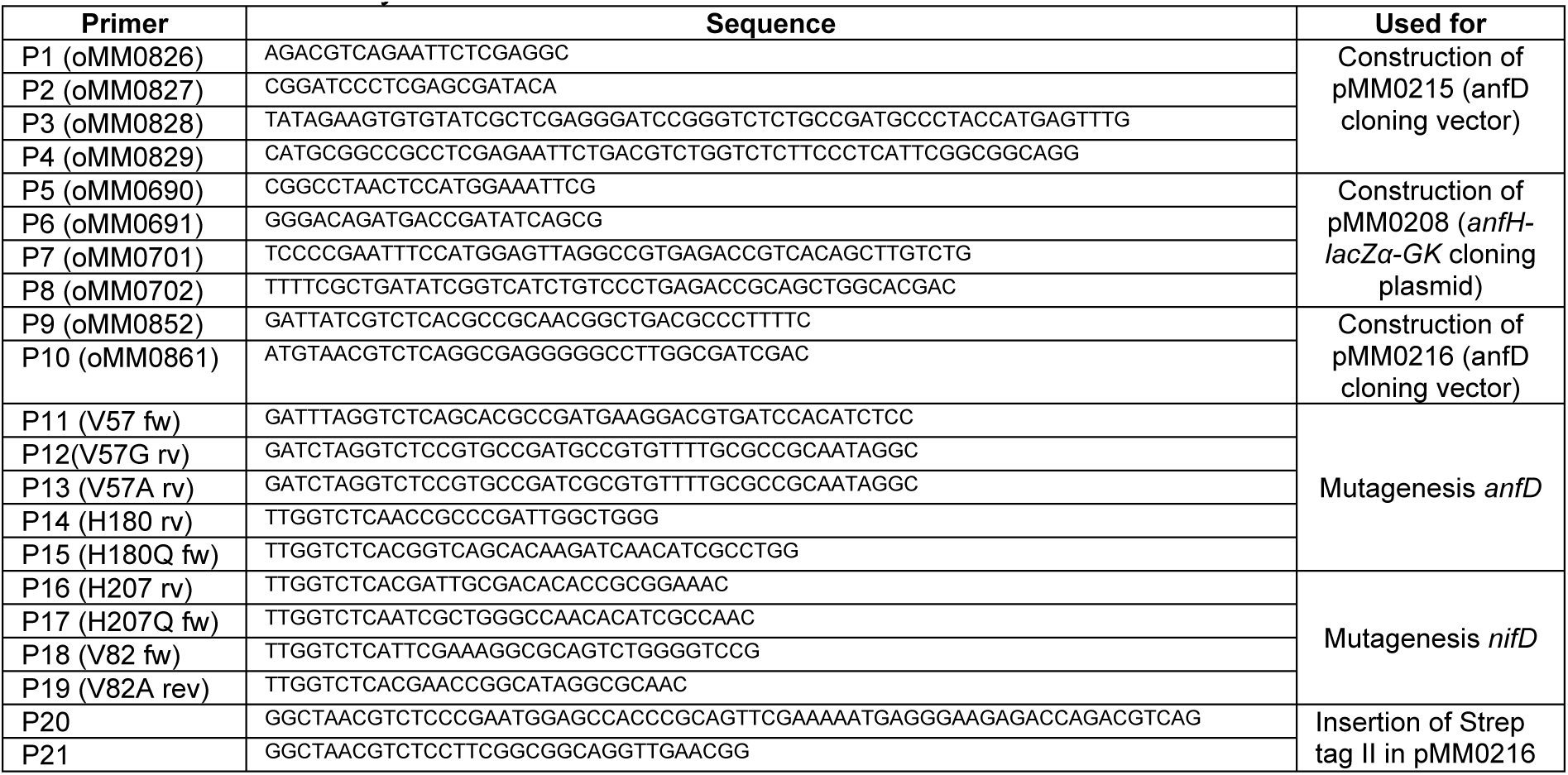
Primers used in this study.

**Table S4.**
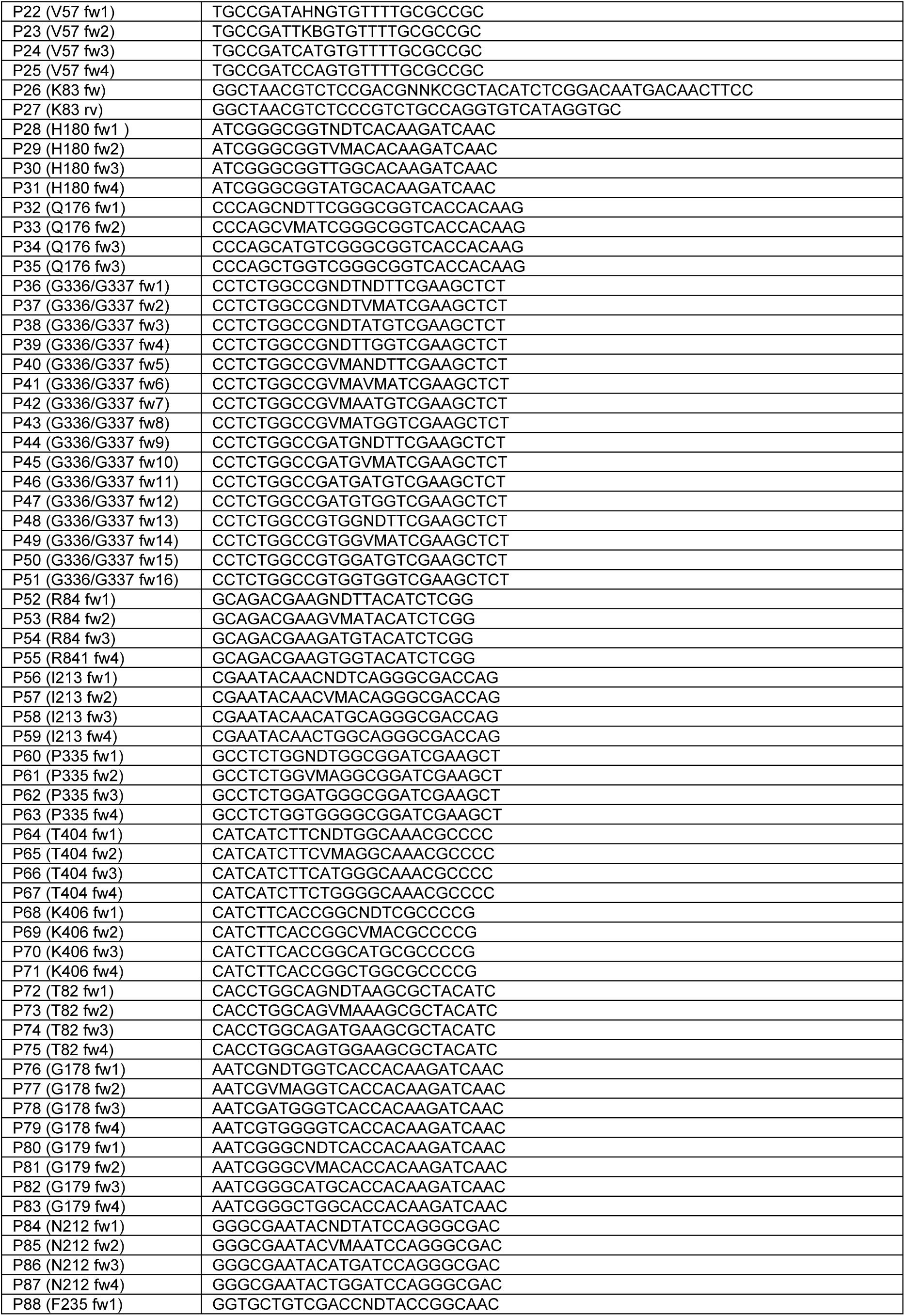
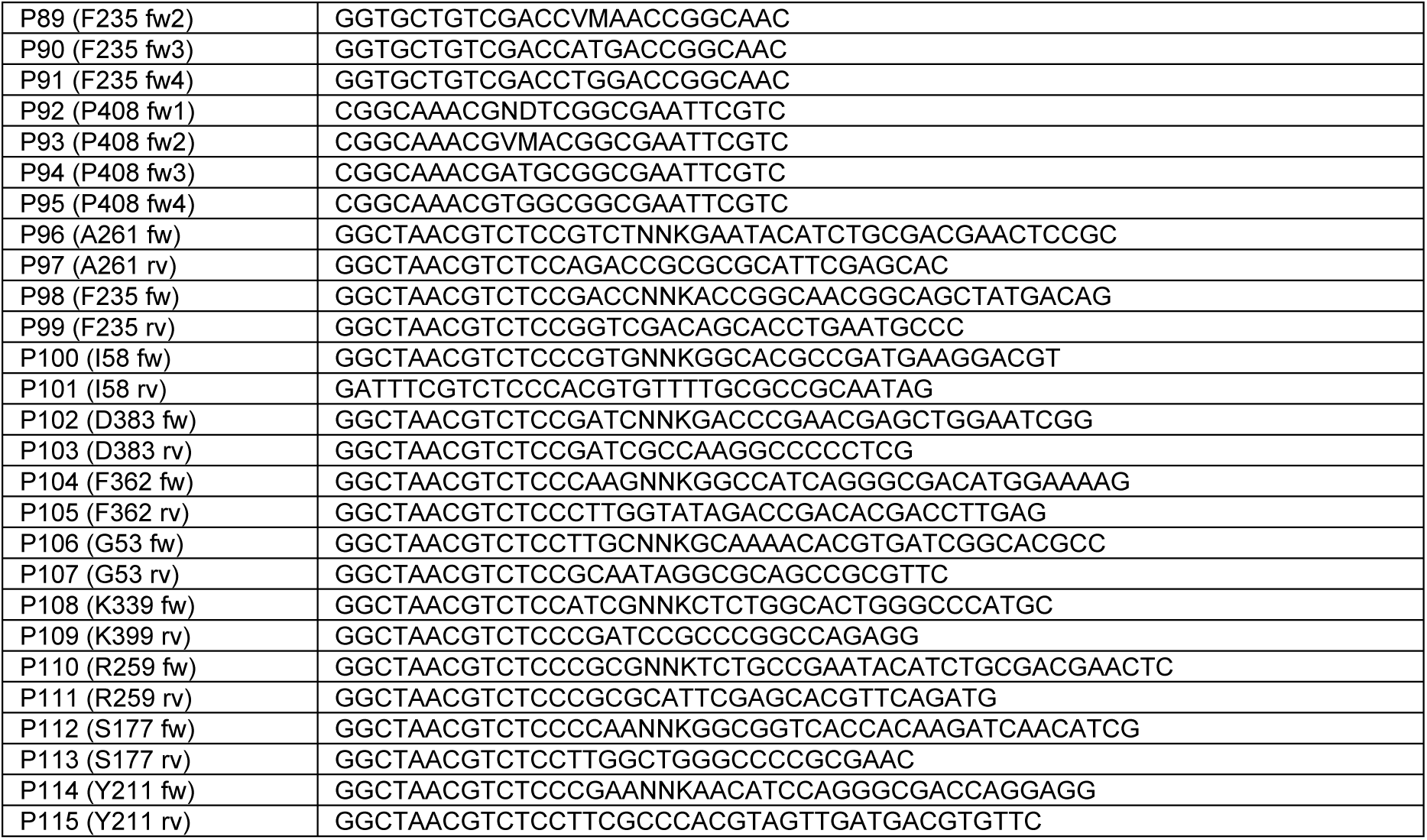
Primers used in library generation of cycle 1.

**Table S5.**
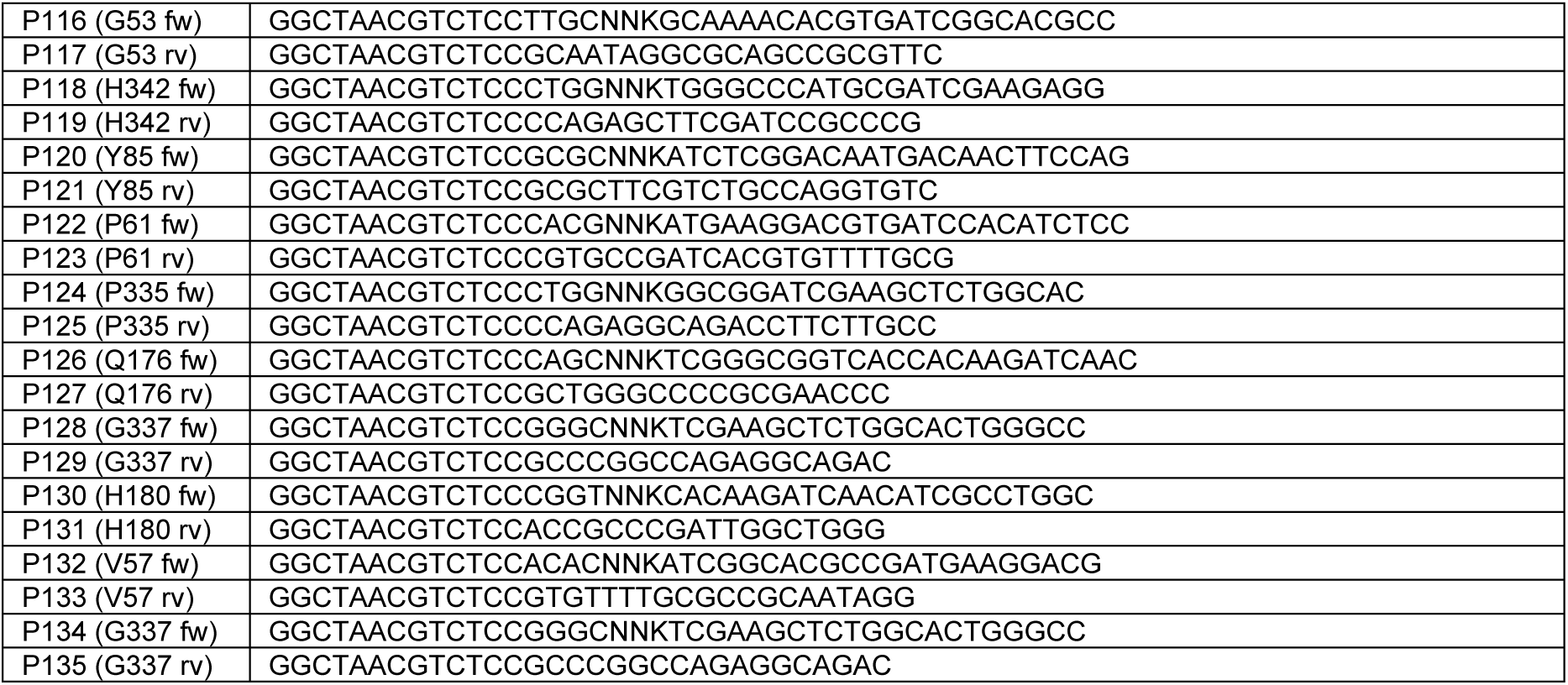
Primers used in library generation of cycle 2.

**Table S6.**
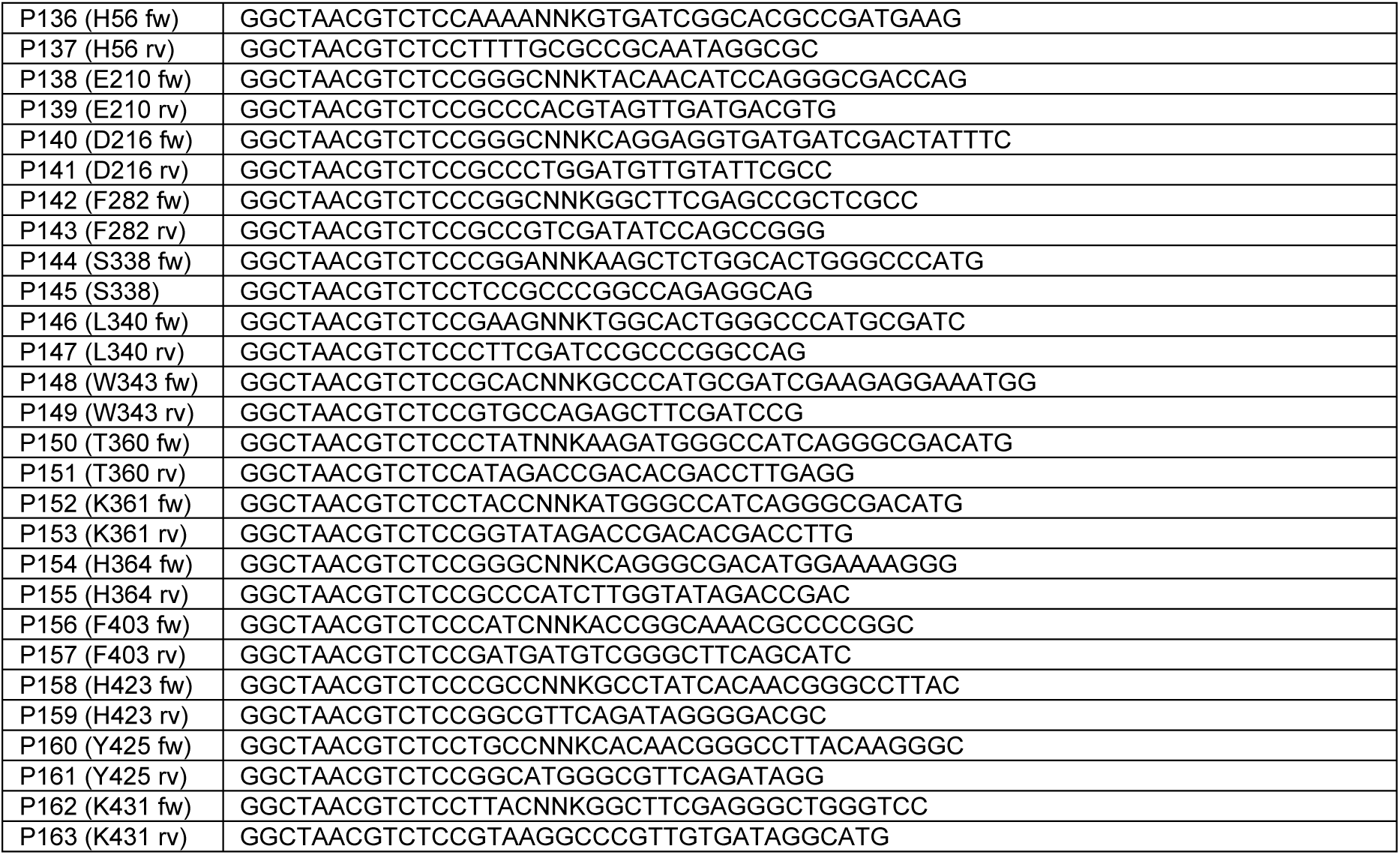
Primers used in library generation of cycle 3.

